# Minimally Invasive DNA-Mediated Photostabilization for Extended Single-Molecule and Super-resolution Imaging

**DOI:** 10.1101/2025.01.08.631860

**Authors:** Michael Scheckenbach, Cindy Close, Julian Bauer, Lennart Grabenhorst, Fiona Cole, Jens Köhler, Siddharth S. Matikonda, Lei Zhang, Thorben Cordes, Martin J. Schnermann, Andreas Herrmann, Philip Tinnefeld, Alan M. Szalai, Viktorija Glembockyte

**Author notes:** These authors contributed equally (Michael Scheckenbach, Cindy Close).

## Abstract

Photobleaching of fluorescence labels poses a major limitation in single-molecule and super-resolution microscopy. Conventional photostabilization methods, such as oxygen removal and addition of high concentrations of photostabilization additives, often require careful fluorophore selection and can disrupt the biological environment. To address these limitations, we developed a modular and minimally invasive photostabilization approach that utilizes DNA-mediated delivery of a photostabilizer directly to the imaging site. Under lower excitation intensities, the DNA-mediated strategy outperformed solution-based approaches, achieving efficient photostabilization at significantly lower additive concentrations. However, at higher excitation intensities, the stability of a single photostabilizer molecule became the limiting factor. To overcome this and reduce the loss of localizations in DNA-PAINT experiments we have also implemented a recovery scheme where the photostabilizer is continuously replenished at the imaging site. We further extended the approach to cell imaging, demonstrating improved localization rate and precision in 3D-DNA PAINT measurements. DNA-mediated photostabilization offers a promising solution for imaging applications where high additive concentrations are prohibited. Its modularity enables adaptation to various imaging schemes and ultimately expands the repertoire of fluorophores suitable for single-molecule and super-resolution imaging.

## Introduction

Single-molecule fluorescence imaging methods have expanded tremendously since the very first observation of single molecules at ultra-low temperatures^1^ and led to many exciting experiments investigating biomolecular interactions, tracking them inside of live cells^2–6^ and even breaking the diffraction barrier to resolve nanoscale features^7^, interactions, or dynamics. Meanwhile one of the main bottlenecks in most of fluorescence imaging experiments remains the premature photobleaching of fluorescent labels^8^. When tracking and monitoring the interactions between individual molecules (e.g. via fluorescence resonance energy transfer (FRET)), the total number of photons that can be collected from a fluorescent label determines the end of the observation window, while in localization-based super resolution imaging techniques it is tightly linked to the localization precision one can achieve^9^.

To extend the total photon budget of fluorescence labels used for these imaging applications one relies on photostabilization strategies that act on the photochemical bleaching pathways (Figure 1a). This typically includes removing molecular oxygen (which can undergo triplet-triplet energy transfer with triplet excited states of fluorescent labels) to prevent the sensitization of singlet oxygen and downstream reactive oxygen species (ROS)^10, 11^. However, removal of oxygen leads to long-lived and reactive triplet dark states, therefore, oxygen removal is typically supplemented by addition of triplet state quenchers (TSQs). ^8, 12^ TSQs can quench the triplet excited states via photophysical mechanisms (k_TET_ in Figure 1a), such as energy transfer (e.g., as is observed for cyclooctatetraene^13–18^ or Ni^2+^ ions^19, 20^) or photochemical mechanisms that rely on reduction (or oxidation) of the triplet excited state with an appropriate reducing (or oxidizing) additive to generate the radical anion (or cation) species^21–24^. The long-lived radical intermediates are subsequently rescued by the addition of a complementary oxidizing (or reducing) partner, an approach that is commonly known as ROXS for **r**educing and **ox**idizing **s**ystem (k_red_ and k_ox_ in Figure 1a)^12^. To circumvent the need for high concentrations of solution-based additives, the photostabilizers can alternatively be directly coupled to the fluorophore core to obtain “self-healing” dyes, however, at the price of additional synthesis and optimization steps.^25–33^ The strategies outlined above have helped to improve the photon budgets by over hundreds of folds in specific instances. Nonetheless, even with the most photostable fluorescent labels paired with the most efficient stabilization approaches the total number of photons is limited to a few millions of photons.

**Figure 1.**
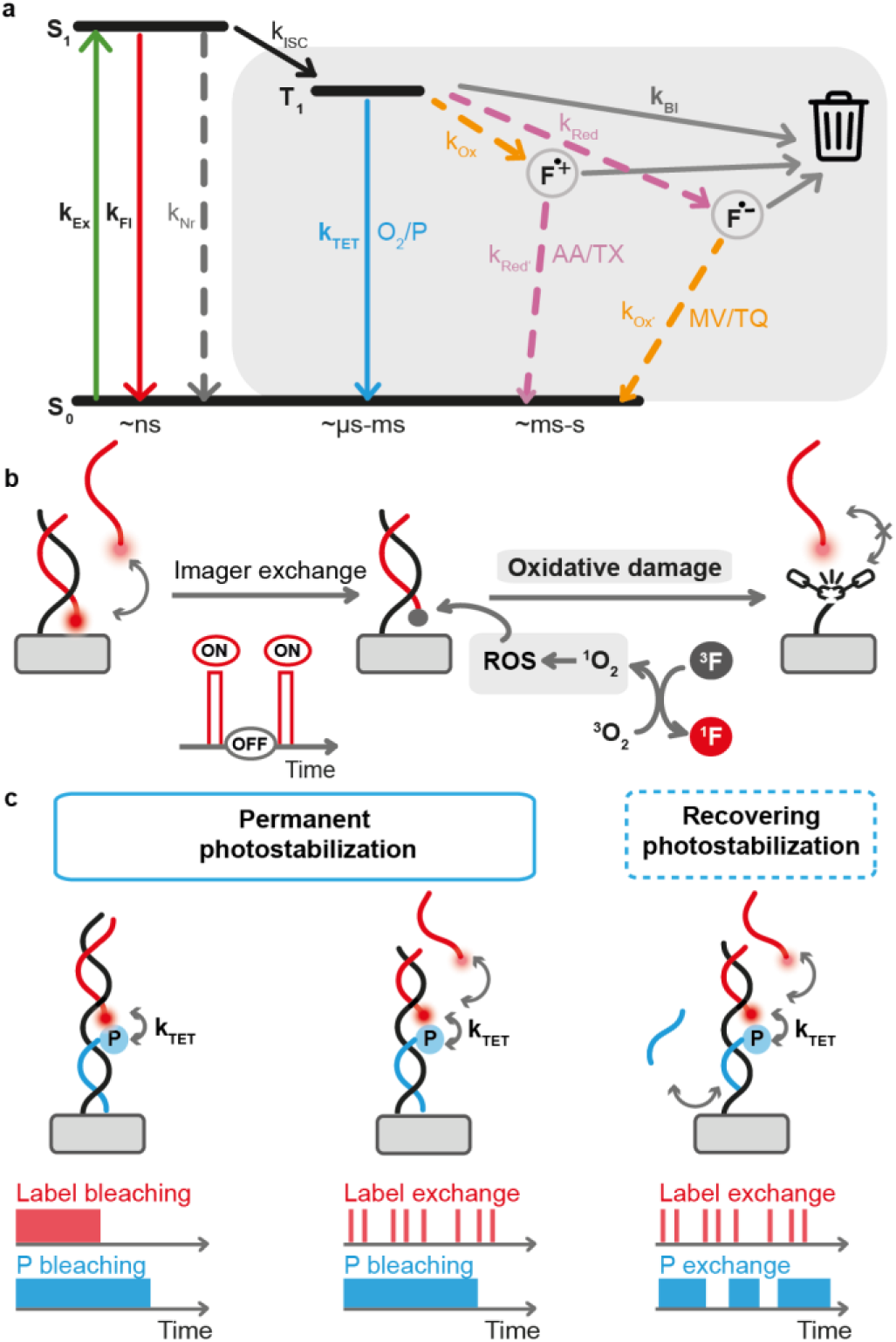
**a)** Jablonski diagram illustrating the photophysical processes involved in photobleaching pathways of fluorescent labels and common strategies to mitigate them by depopulating the non-emissive and reactive triplet and radical states. Here one can utilize photophysical triplet state quenchers that operate via triplet energy transfer (TET, k_TET_) or photochemical quenchers that rely on ping-pong redox reactions (ROXS, k_red_, k_ox_); **b)** Illustration of oxidative damage that limits the performance of imaging methods based on continuous label exchange: although the photobleached label is exchanged the photostabilization is necessary to ensure the depopulation of the reactive triplet states and generation of ROS and consecutive photodamage of the docking site; **c)** Minimally invasive photostabilization introduced in this work that relies on DNA-mediated delivery of the photostabilizer directly to the imaging site. The photostabilizer can be attached to the imaging/docking site permanently (left and middle panels) via stable DNA-DNA interaction or exchanged continuously using transient DNA-DNA interaction (right panel). Lower panel illustrates time course of the experiment and residence time of the fluorescent label (red) as well as the photostabilizer (blue) on the docking site.

One of the imaging “tricks” used to fundamentally overcome the limits posed by the finite photon budget of single fluorescent labels involves the continuous replacement of bleached labels via transient binding. This is nicely exemplified by super-resolution imaging with DNA-PAINT.^34^ DNA-PAINT relies on transient binding of short fluorescently labelled DNA oligonucleotides (imager strands) to the target of interest labelled with a complementary DNA sequence (docking sites) to achieve apparent blinking at the imaging site which is, in turn, used for stochastic super-resolution imaging. Here, each docking site can bind multiple imager strands over time and the imaging quality and efficiency are no longer limited by the photobleaching of the single-fluorescent label in contrast to other localization-based super-resolution imaging methods (e.g., PALM or STORM)^35–37^.

In recent years, several other approaches that exploit DNA-mediated dynamic exchange of fluorescent labels to generate a long lasting and photostable fluorescence signal have been put forward. For example, in our previous work we have used the dynamic exchange of bleached fluorophore strands with intact ones from solution to generate self-regenerating DNA origami-based brightness rulers.^38^ Repetitive DNA binding motives to continuously exchange labels in solution have also been successfully exploited to design a long-lasting fluorescence label for tracking single biological molecules for hours.^39^ Introduction of DNA-PAINT imager strands into STED microscopy has overcome the photobleaching of permanent fluorescent labels.^40^ The more recent “REFRESH” and “Dye cycling” approaches used analogous strategies to continuously exchange both, donor and acceptor labels enabling near-continuous observation of single-molecules for more than an hour and extending these ideas to FRET imaging studies.^41, 42^ New imaging schemes going beyond DNA-mediated transient binding have also been realized, e.g. by engineering exchangeable HaloTag ligands that can be used for super-resolution imaging.^43, 44^

These dynamic labelling strategies elegantly overcome the problem of bleaching of single fluorescent labels. However, they are still limited by the photochemical processes in the excited states. When imaging is performed in the absence of photostabilization additives, every time the fluorophore enters the triplet excited state it has a probability to generate singlet oxygen and other ROS. While the ROS-induced photodamage to the fluorescent label is addressed by recovering it over time, the damage to the target molecule or binding site is not mitigated (Figure 1b). For example, in DNA-PAINT imaging studies it has been shown that the photoinduced damage of docking sites leads to loss of localizations over time, setting a limit on the total number of localizations that can be achieved.^45^ Not surprisingly, removal of oxygen and use of common photostabilization cocktails, such as ROXS, has also been essential in the above-mentioned studies using DNA-mediated label exchange^38, 39, 41, 42^, emphasizing the importance of photostabilization even if the experiment is no longer limited by the bleaching of the label itself. ^38, 39, 41, 42, 45^

Nevertheless, the need for oxygen removal and addition of photostabilizing agents also limits their use to applications compatible with the required conditions. On one hand, photostabilization additives at millimolar concentrations can influence the biomolecular system under study^46^ and the removal of oxygen by enzymatic scavenging systems can result in acidification of the sample solution^47, 48^. On the other hand, efficient removal of oxygen and solution-based photostabilization, which depend on efficient diffusional collision, simply might not be possible (e.g. crowded and inaccessible cellular compartments^49, 50^, correlative measurements^51^). Additionally, in applications that rely on the exchange of fluorescent labels in solution such as DNA-PAINT or dynamic labeling, solution-based photostabilization can lead to an undesirably high background (unspecific photostabilization). In multi-color imaging schemes, it can also be difficult to identify one photostabilization additive that allows for optimal performance of multiple fluorophores.^20^

With these limitations in mind, we developed a modular and minimally invasive photostabilization strategy which relies on the DNA-mediated delivery of a photostabilizer, i.e. a TSQ, directly to the imaging site, circumventing the need for high concentrations of additives. We first characterize and benchmark this strategy by comparing it to solution-based photostabilization and then show that it can be successfully applied to slow down photoinduced depletion of docking sites in DNA-PAINT imaging as well as utilized for long term imaging studies based on continuous exchange of labels. Finally, we use this photostabilization strategy to enable 3D DNA-PAINT imaging in cells in the presence of oxygen to mimic imaging in biological samples, where oxygen removal is not feasible. To illustrate the future modularity of this approach, we also outline how it can be extended to different imaging schemes.

## Results

In DNA-PAINT, the target structure is chemically modified with a nucleic acid sequence. To not only direct the imager strand but also the photostabilizer to the imaging site, we extended the DNA docking site sequence. The additional binding site for the photostabilizer strand allows it to locally act at the imaging site, where most of the photoinduced damage occurs (Figure 1c). By changing the sequence length one can design the photostabilizer strand to either permanently bind at the imaging site (Figure 1c, left and middle panels) or continuously exchange analogously to fluorescent labels (Figure 1c, right panel). Differently from a “self-healing” approach which requires direct coupling of the photostabilizer to the fluorophore, our strategy relies on coupling the photostabilizer to a DNA oligonucleotide which can later be modularly reused for different fluorescent labels or imaging schemes. To avoid potential radical intermediates, in this work, we used cyclooctatetraene (COT) as physical TSQ due to its ability to depopulate the triplet excited states via a photophysical pathway circumventing possible radical intermediates at the imaging site. COT-functionalized oligonucleotide photostabilizers were prepared using a previously reported universal linker molecule by coupling maleimide functionalized COT linker molecules to thiolated DNA oligonucleotides (Scheme S1, Table S4).^52^

We first performed single-molecule fluorescence studies to test whether DNA-mediated photostabilization can be as efficient as solution-based photostabilization. Common additives work at millimolar concentrations, thereby proving a virtually unlimited pool of photostabilizer molecules. To show the strength of our photostabilization approach we chose the otherwise photolabile Cy5 dye. As it has been demonstrated that COT significantly improves photostability of Cy5^30^, we expected a distinct contrast between the bare and photostabilized fluorophore. For this, Cy5-labelled twelve helix bundle DNA origamis (12HB) were immobilized on a BSA-biotin passivated glass coverslip using neutravidin-biotin interactions (Figure S2, Figure 2) and imaged on a total internal reflection (TIRF) microscope. The COT photostabilizer strand (17 nucleotides long) was permanently attached to the imaging site via DNA-hybridization (pCOT, Figure 2c). Control samples included a construct carrying an analogous oligonucleotide without the COT moiety (Figures 2a and 2b). In the absence of oxygen (scavenged with glucose oxidase/catalase) and photostabilization additives, single-molecule imaging of Cy5 resulted in a characteristic fluorescence blinking behavior due to the formation of long-lived triplet-born dark states (Figure 2a).^12, 17, 20, 24^ This dark state formation also leads to early saturation of fluorescence signal at excitation intensities as low as 0.3 kW/cm^2^ (Figure 2a and 2d). Subsequent addition of 2 mM of triplet state quencher COT^53, 54^ allowed for efficient collisional quenching of triplet excited states, consequently leading to a much more stable and bright fluorescence signal (Figure 2b and 2d).

**Figure 2.**
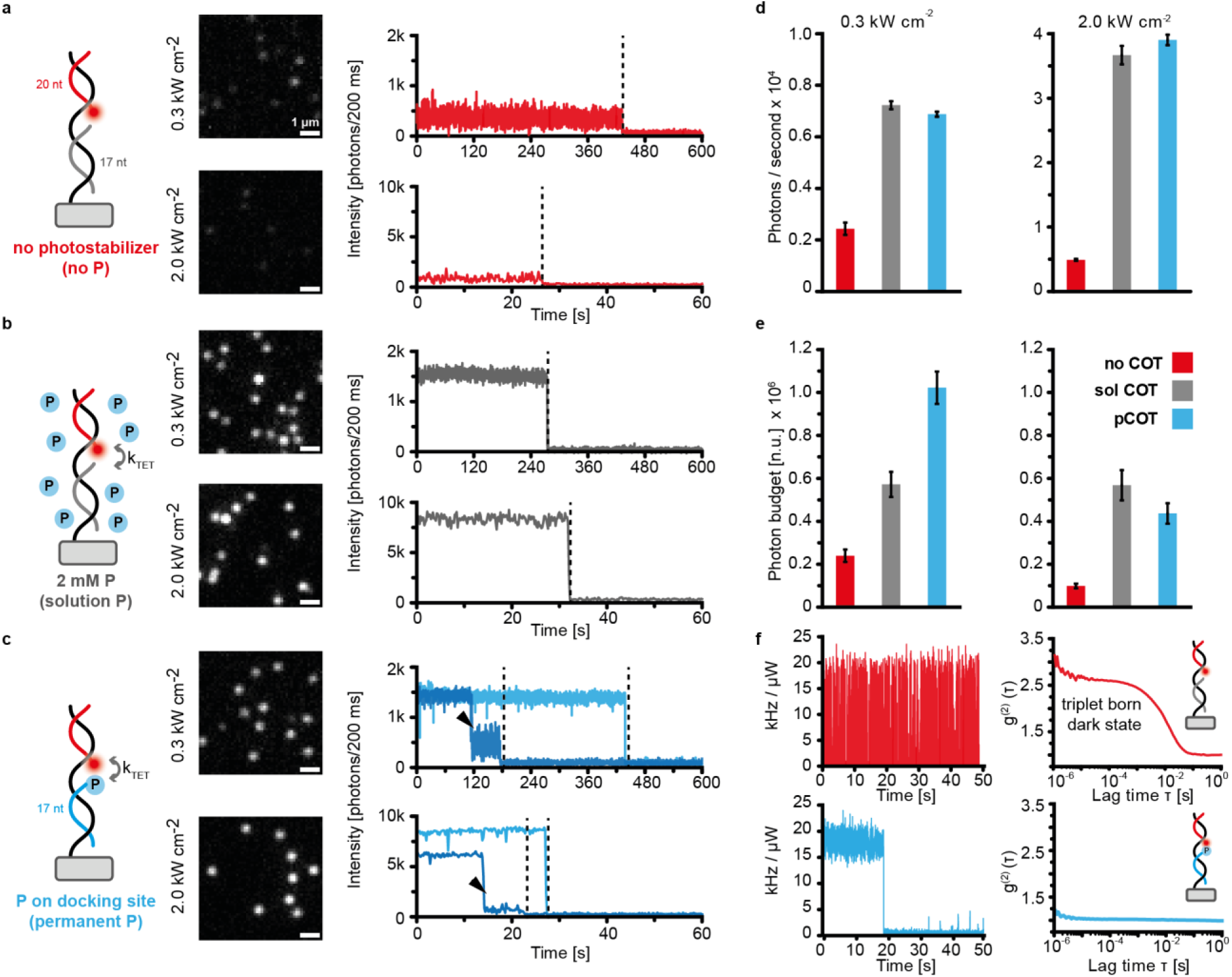
Photophysical characterization of the DNA-mediated photostabilization strategy. Single molecule TIRF images (middle panel) and representative single-molecule trajectories (right panel) obtained for Cy5-labelled DNA origami in the absence of oxygen and (**a**) no photostabilization additives, (**b**) 2 mM COT as a photostabilizer, and (**c**) DNA-labelled COT photostabilizer attached directly at the imaging site; **d)** Average brightness of single-molecule fluorescence signal and (**e**) average total photon budget obtained at two different illumination intensities; **f)** representative single-molecule trajectories obtained imaging the Cy5B-labelled DNA origami in the absence of oxygen and no photostabilization additives (top) or in the presence of DNA-mediated photostabilization by COT (bottom). The corresponding average fluorescence autocorrelation functions obtained analyzing single molecule trajectories are shown on the right and indicate efficient DNA-COT mediated depopulation of the dark states.

Aiming for efficient collisional quenching between the photostabilizer (COT) and the fluorophore (Cy5) in the DNA-mediated strategy^14^, we designed the DNA docking site in a manner that leads to a head-to-head placement of the two (Figure 2c)^17^. As illustrated in the single-molecule fluorescence trajectories (Figure 2c), a single photostabilizer delivered to the imaging site via DNA hybridization was sufficient to achieve a stable fluorescence signal, as bright as the one obtained in the presence of 2 mM COT as a solution additive (Figure 2d). In fact, further characterization of total photon budget (average number of photons collected before the photobleaching event) demonstrated that at low illumination intensities (0.3 kW/cm^2^, typical for single-molecule studies) a DNA-mediated approach is more efficient leading to an almost two-fold higher photon budget when compared to the solution-based approach (Figure 2e). This suggests that the direct delivery of photostabilizers to the imaging site with the help of DNA hybridization, resulting in higher local concentration, can be even more efficient than collisional quenching by solution additives.

To investigate the effectiveness of the DNA-mediated photostabilization approach for applications that require higher illumination intensities (e.g., single-molecule localization microscopy (SMLM)) we also carried out single molecule studies at 2.0 kW/cm^2^. Under these conditions, however, the photostabilization with a single COT moiety resulted in lower overall photon budget when compared to 2 mM COT in solution. We hypothesize that this reduced performance could be related to photoinduced degradation of the COT moiety at increased excitation intensities. In par with this observation, single-molecule fluorescence trajectories for both a low and a high excitation power density revealed instances of Cy5 fluorescence blinking before bleaching (9% of the traces for 0.3 kW/cm^2^, 15% of the traces for 2.0 kW/cm^2^, Figures S3 and S4). This observation additionally illustrates that the stability of the photostabilizer itself can present a bottle-neck in the performance, especially in the photostabilization schemes that rely on only one photostabilizer moiety, i.e. as in the one studied here or in self-healing dyes. ^30^

To confirm that the improved photostability stems from efficient depopulation of the triplet excited states and shed light on efficiency of triplet state quenching via the DNA-mediated approach, we performed analogous single-molecule imaging studies with the rigidified Cy5 analogue Cy5B.^55, 56^ For this dye, a dark state originating from photoisomerization can be excluded. Autocorrelation analysis of single-molecule fluorescence trajectories of Cy5B in the absence of oxygen revealed analogous blinking due to the formation of triplet-born dark states (Figure 2f, upper panel). Photostabilization via DNA-mediated strategy with pCOT, on the other hand, led to a stable and bright fluorescence signal with significant quenching of the triplet-born dark state intermediates, confirming an efficient collision between the fluorophore and the photostabilizer.

The limitation posed by having only a single photostabilizer that can eventually photodegrade is even more pronounced in imaging applications that rely on the continuous exchange of fluorescent labels under continuous illumination, such as DNA-PAINT^30, 34, 45^ and recovering labeling.^38, 39, 41, 42^ Over the course of an experiment, the single photostabilizer molecule has to stabilize multiple fluorophores binding transiently over time, favorably under a high excitation illumination to ensure a high photon count rate. To investigate the applicability and performance of DNA-mediated photostabilization under these conditions, we implemented a shorter binding sequence (11 nucleotides) for the transient, recovering binding of Cy5 imagers and compared it to the permanent labeling approach (Figures 3, S8-10).^41^ We performed long-term single molecule studies over an observation time of 60 min at low illumination intensities of 0.1 kW/cm^2^ to ensure that photobleaching is slow and not outcompeting transient binding kinetics. Even though efficiently photostabilized by a single COT moiety on the DNA docking site, the permanent Cy5 label (Figure 3a) was still limited by irreversible photobleaching yielding an average total photon budget of around 1×10^6^ photons (Figure 3d) comparable to the one obtained for slightly higher illumination intensity used before (0.3 kW/cm^2^, Figure 2e). Nevertheless, as observed earlier, the DNA mediated photostabilization approach with pCOT again outperformed the commonly used solution photostabilization with 2 mM COT by a factor of ca. 2, while yielding similar photon count rates (Figure 3d blue vs. grey bars, more data in Figures S8 and S9).

**Figure 3.**
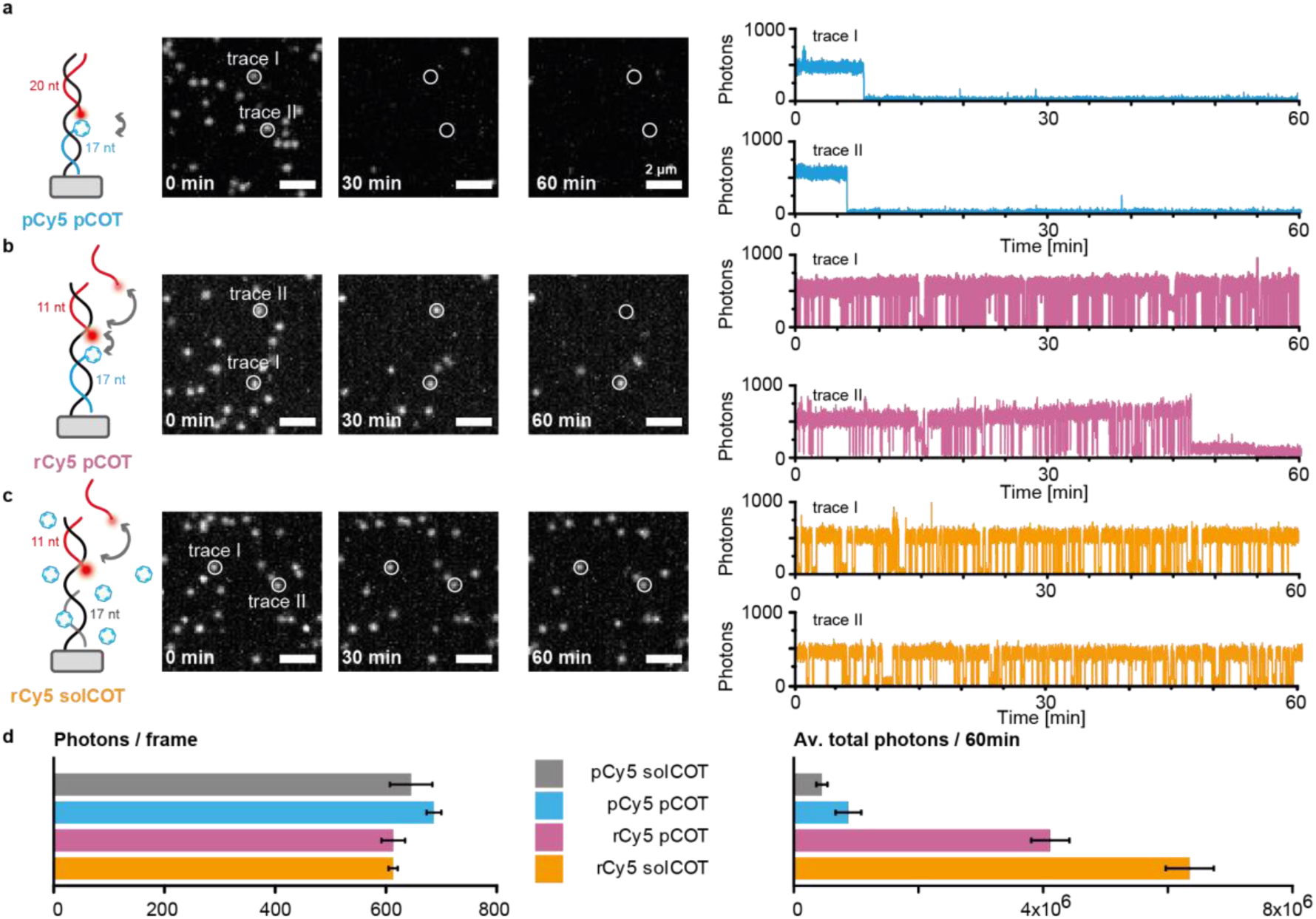
DNA-mediated photostabilization for recovering Cy5 imager labels in the absence of oxygen over a long observation time (60 min) and under low illumination intensity (ca 0.1 kW/cm^2^). Single molecule TIRF images at 0, 30, and 60 min as well as representative single-molecule trajectories obtained for (**a**) a permanent Cy5 and a pCOT on the DNA docking site, (**b**) a recovering Cy5 and a permanent COT on the DNA docking site, and (**c**) a recovering Cy5 label with 2 mM COT as a photostabilizer in solution, respectively; **d**) Average brightness of single-molecule fluorescence signals and average total photon budget obtained for different imaging conditions. Bar plots in (**d**) represent average of three measurements, errors represent the standard deviation.

Switching to a dynamic imaging scheme and using 10 nM of the recovering Cy5 imager (Figure 3b and S10) came at the cost of a slightly increased background but resulted in blinking, pseudo-continuous trajectories (as observed in previous studies^41, 42^) with comparable brightness values, but a highly improved imaging time, surpassing the total photon budget of a single Cy5 molecule by approximately four-fold (Figure 3d and S10). However, we still observed slow photo-induced degradation of the DNA docking site over time, leading to a loss of around 60% of imaging trajectories after 60 min (Figure S11). Recovering imaging with solution based photostabilization (Figure 3c), resulted in an even higher photon budget (ca. 6×10^6^ photons over 60 min) and almost no label bleaching over the entire duration of the experiment (Figures 3d, and S11). On one hand, these findings highlight that DNA mediated photostabilization can be applied to recovering labels resulting in long-lasting single-molecule observation times breaking the photobleaching limit of a single fluorophore. On the other hand, they also underscore that even under very low illumination intensities, photoinduced damage to the photostabilizer remains the bottleneck, especially when the experiment is no longer limited by the bleaching of the fluorescence label and requires long-observation times.

After successfully applying DNA mediated photostabilization to recovering imager labels under low illumination intensities applicable for single-molecule imaging routines such as single-particle tracking or SM-FRET studies, we next aimed to extend our approach to DNA-PAINT super-resolution imaging. In SMLM techniques such as STORM, PALM or DNA-PAINT, the achieved resolution in the super-resolved image relies on the photon count rate of the detected blinking events. SMLM experiments are, hence, commonly performed under high illumination intensities (typically ≥ 1.0 kW/cm^2^) to obtain a high spatiotemporal resolution. To test the performance and stability of DNA mediated photostabilization under these conditions, we equipped a 12HB DNA origami with three DNA docking sites placed at 90 nm distances (Figure 4). Each docking site consisted of a short DNA-PAINT imager binding sequence (8 nucleotides) and a neighbouring photostabilizer binding sequence of different lengths in order to investigate permanent (17 nt for pCOT) as well as recovering (10 nt for recovering COT (rCOT)) photostabilization schemes (Figure 1).

**Figure 4:**
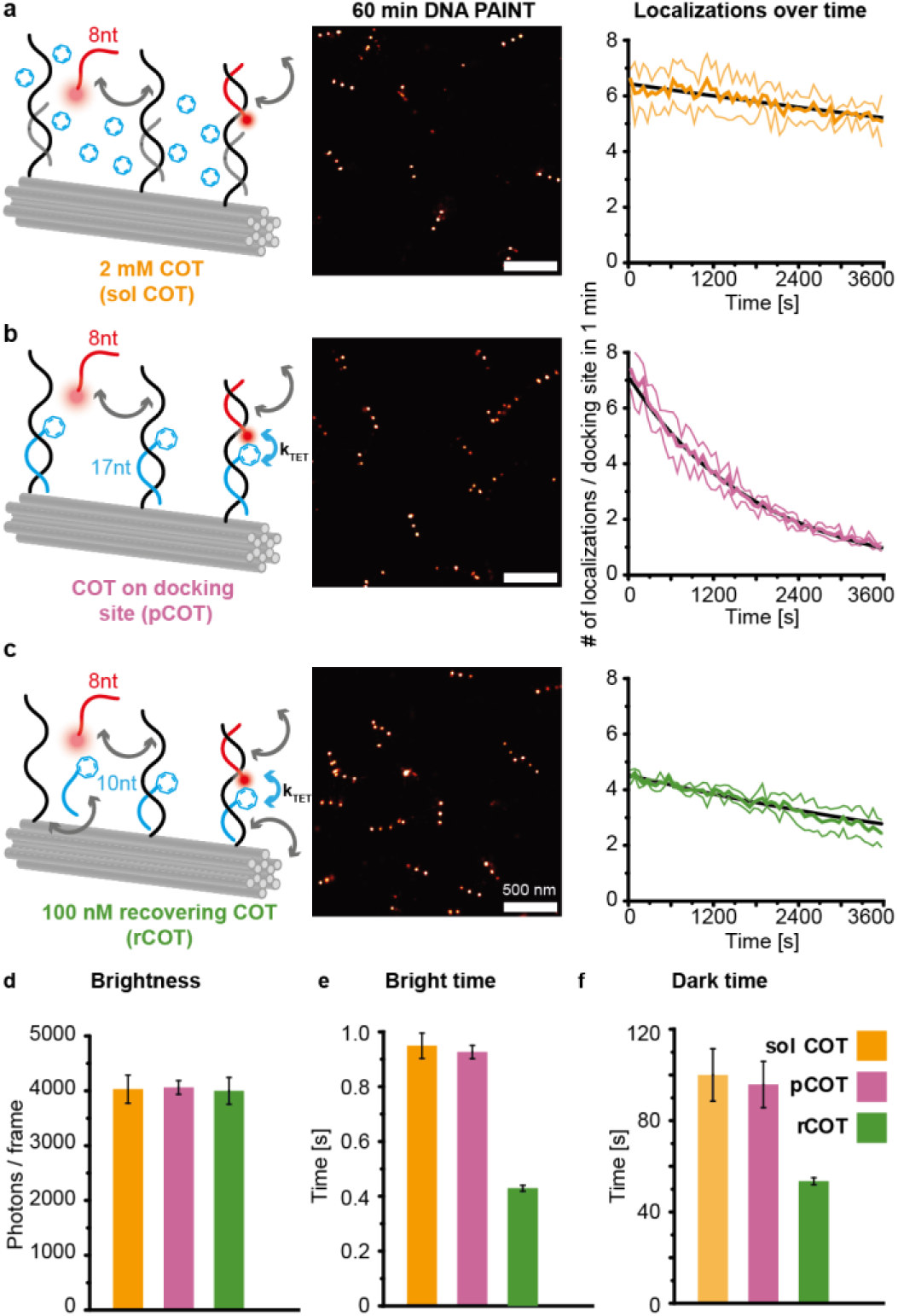
DNA-mediated photostabilization for DNA-PAINT imaging with Cy5 in the absence of oxygen and under high illumination intensity (ca 1.0 kW/cm^2^). Exemplary reconstructed DNA-PAINT images of 12HB nanostructures with three docking sites and detected localizations per docking site over time obtained with (**a**) 2 mM COT solution-based photostabilization, (**b**) a permanent DNA mediated photostabilization (pCOT, 17 nt binding sequence) and (**c**) a recovering DNA mediated photostabilization (rCOT, 10 nt binding sequence). Brightness values (**d**), and DNA-PAINT bright times (**e**) and dark times (**f**) extracted for single DNA docking sites. Coloured curves in (**a**) to (**c)** represent the average of three measurements, errors represent the standard deviation, dark lines represent exponential fits. Bar plots in (**d**) to (**f**) represent the average of three measurements, errors represent the standard deviations.

For this purpose, we performed DNA-PAINT imaging on a total internal reflection (TIRF) microscope with 1 nM of a Cy5 imager strand in an oxygen depleted imaging buffer at a typical SMLM illumination intensity of 1.0 kW/cm^2^. DNA-PAINT imaging in oxygen-depleted buffer without any photostabilization additives resulted in a poorly resolved image and localizations bearing low photon counts (660 photons/ 100 ms) at a generally low localization rate over time indicating that the presence of TSQ is crucial for successful super-resolution measurement (Figure S12). DNA-PAINT measurements with the classical solution-based stabilization (2 mM COT, Figure 4a), on the other hand, allowed for the successful reconstruction of the designed three-spot pattern enabling the selection and examination of individual docking sites. To investigate the stability of the DNA docking sites against triplet state mediated and ROS-induced photodamage, we extracted the number of localizations of individual docking sites for a defined time unit (i.e., 1 min). As reported previously, the addition of an unlimited pool of photostabilizer molecules in solution resulted in an almost constant average localization rate per single docking site over the whole observation time indicating a high stability of the DNA docking sites (Figure 4a, right).^45^ Next, we performed DNA-PAINT imaging on the 12HB nanostructures with a COT functionalized photostabilizer strand permanently bound to the docking site (pCOT, Figure 4b). While the reconstructed DNA-PAINT images revealed the designed three-spot pattern accurately and photon count rates similar to solution-based photostabilization, we also observed a rapid decay of localizations over time and almost complete loss of docking sites by the end of the 60 min measurement (Figure 4b, right). Due to the high turnover of imager strands and high illumination intensity, the limited stability of the COT photostabilizer became even more relevant limiting the meaningful observation times to less than 60 min and potentially preventing a complete reconstruction of the sample.

Although the intrinsic instability of COT towards photoinduced electron transfer reactions with oxygen can be improved by the introduction of electron withdrawing groups, this strategy only slows down the irreversible degradation of the photostabilizing molecule and does not overcome it entirely.^30, 57^ To circumvent the inevitable loss of docking sites due to limited stability of the COT moiety, we introduced an imaging scheme that allows for the recovery of the photostabilizer strand as well (rCOT, Figure 1c), analogously to the recovery of bleached fluorescent labels. To this end, we shortened the binding sequence of the photostabilizer strand on the DNA docking to 10 nt to ensure a shorter binding time and dynamic exchange. After determining the concentration of the recovering COT strand needed to saturate the binding to the DNA docking site (Figure S13), we then carried out the recovering DNA-PAINT photostabilization with 100 nM rCOT photostabilizer over 60 min (Figure 4c). Photon count rates comparable to pCOT and solution-based photostabilization (Figure 4d) and reconstruction of the DNA-PAINT images revealing the designed three-spot pattern suggested an efficient photostabilization for the rCOT strategy despite its dynamic nature. Moreover, the average localization rates per single docking site showed a significantly improved stability of the DNA docking sites when compared to pCOT photostabilization which relies on a single photostabilizer. Therefore, by relying on this recovering exchange of photostabilizer molecules with the help of DNA, we could achieve photostabilization comparable to the solution-based approach, however, at seven orders of magnitude lower concentration of additives (Figure 4c, right), also in an oxygenated environment (Figure S14).

To determine if the binding kinetics of the DNA-PAINT imager are affected by the COT functionalization on the docking site, we also extracted the average dark- and bright-times of selected docking sites for each photostabilization approach (Figures 4e, 4f and S15). While we found comparable binding kinetics for the solution-based approach and the pCOT photostabilization, the dynamic rCOT photostabilization surprisingly resulted in the decrease of both, bright-times and dark-times, by ca. 50%, in turn, doubling both the association and dissociation rates of the imager strand to the DNA docking site. The faster blinking for the sample in the rCOT imaging scheme was also clearly visible in fluorescence time traces (Figure S16), highlighting the increased binding and dissociation rates.

To explore whether DNA-mediated photostabilization could be used for imaging applications with more complex biological samples, we performed DNA-PAINT measurements in fixed fibroblast cells (COS-7) using the pCOT photostabilization strategy. We chose to image microtubules, as they are an established model system in the super-resolution microscopy community, allowing for an intuitive and fair comparison to different labeling and photostabilization techniques. Experiments were performed in ambient oxygen conditions, to mimic applications where the use of oxygen scavenging systems is prohibited. This way, COT bound to the docking site directly competes with the high concentrations of oxygen in solution (typically ca. 0.3 mM).^11^ To observe the effect of pCOT stabilization on the performance of Cy5, we imaged the microtubules for 60 minutes (Figure 5a). Figure 5b shows zoom-ins of reconstructed images from both conditions with and without DNA mediated photostabilization. Homogenous illumination of the sample was ensured by including a flat-top beam shaper in the excitation path.^59^ The photon number per localization is color coded, illustrating the increased number of photons for DNA-PAINT imaging with pCOT photostabilization. To quantify this further, we plotted the binned number of localizations over the course of the experiment (Figure 5c). The inset of Figure 5c shows how, during the experiment, localizations continuously increase in the entire field of view, while the pCOT sample has an overall higher number of localizations to begin with and accumulates them more quickly. This trend is confirmed when comparing selected regions of interest (ROIs) of fixed dimensions within several individual microtubules. We hypothesize that the lower number of localizations is either due to the loss of docking sites when Cy5 is not photostabilized or an effect of Cy5 bleaching within the binding time leading to an insufficient number of photons per localization to be detected. With COT, an increased fraction of docking sites is preserved, leading to the higher number of localizations.

**Figure 5.**
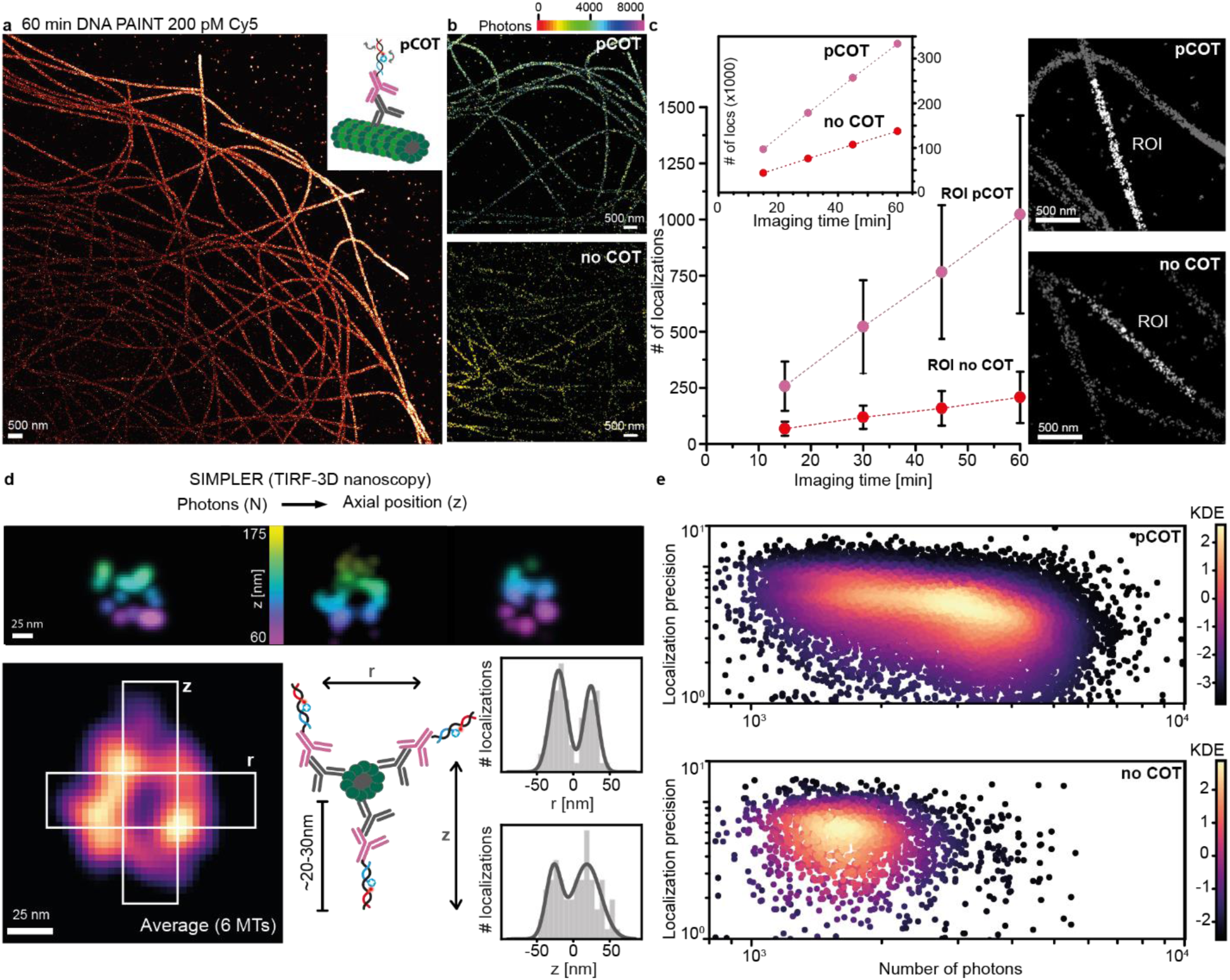
Application of DNA-mediated photostabilization to DNA-PAINT imaging in cells. 60-minute DNA-PAINT experiment under low illumination intensities (ca 0.6 kW/cm^2^) and with ambient oxygen. **a)** Overview image of the sample where the permanent photostabilizer was added at 200 pM (inset shows the labelling strategy); **b)** Exemplary zoom-ins on regions in the samples with no photostabilization or pCOT photostabilization (color coded by photon number); **c)** Accumulation of localizations over the course of the experiment within comparable selected regions (example highlighted in ROI in white) for pCOT and no photostabilization control. Inset shows localizations over time for the entire imaged region (isolated region of interest (ROI) at a similar position in the field of view for both samples). **d)** Cross-sections of microtubules extracted using the SIMPLER (supercritical illumination microscopy photometric z-localization with enhanced resolution) algorithm.^58^ Three exemplary reconstructed images of microtubules, color coded by position in z-dimension, (top), average of six cross-sections (bottom left), linkage-errors due to labelling reported in literature, histogram of localizations in dimension r and z of the average microtubule (bottom right). **e)** Number of photons plotted against the corresponding localization precision for pCOT and no photostabilization (color coded by kernel density estimation).

To test whether DNA-PAINT measurements with the photostabilized Cy5 can be used to reconstruct the three-dimensional position of localizations, we applied the SIMPLER (supercritical illumination microscopy photometric z-localization with enhanced resolution) algorithm.^58^ The method by Szalai *et al.* converts the number of detected photons to the axial position (z) of single molecules when acquisitions are performed under total internal reflection (TIR) conditions. SIMPLER strongly relies on the use of stably emitting dyes, since fluctuations in brightness significantly reduce axial localization precision. Given that a binding event needs to last at least three camera frames to be considered in the SIMPLER algorithm, it is crucial that the dye does not undergo fast photobleaching once the imager strand binds to the docking site. Figure 5d shows cross-sections of three exemplary microtubules, as well as an average over six microtubules. Considering the size of the primary and secondary antibodies (adding approximately 20-30 nm^60^) the achieved peak-to-peak distance of the hollow microtubule (44 nm in r and 45 nm in z) is in good agreement with literature.^61, 62^ Inherently, the localization precision in DNA-PAINT measurements is a function of the photon number N (Figure 5e). Additional to yielding overall more localizations, the introduction of DNA-mediated photostabilization increases the number of localizations with higher photon count. The mean photon count for pCOT amounts to 3082, while without COT this value drops to 1897. As a result, the ratio of localization precision pCOT/no COT (σ_x_ = 5.6 nm/6.0 nm) is 0.91 (ratio of √*N* = 0.76).

To investigate whether even better performance can be achieved we also performed experiments with the recovering rCOT strand in the imaging solution (Figure S18). However, no substantial improvement of brightness or number of localizations was observed when compared to the pCOT stabilization shown in Figure 5. Since super-resolution experiments in cells were performed at lower excitation intensities (0.6 kW/cm^2^), we hypothesize that under this regime we were not limited by bleaching of the photostabilizer as observed previously (Figure 4). Together, this illustrates, that the choice between permanent or recovering modality can be based either on sample requirements (e.g., when pCOT would be the least invasive choice) or imaging conditions (e.g., when higher illumination intensity is necessary, rCOT can potentially help to circumvent photostabilizer bleaching).

## Discussion

Using DNA interactions to direct the photostabilizer to the imaging site enables the modular combination of a photostabilizer with different fluorophores (Figure 6a, first row). Within this work, we attached COT to oligonucleotides of two different lengths (10 and 17 nt) and applied them to various fluorophores and imager strands on the DNA docking site. While we could show highly efficient photostabilization for permanent and recovering imager strands with the red-emissive dyes Cy5, Cy5B and Atto647N (Figure S7), we observed no photostabilization for permanent Cy3 (Figure S5) and Cy3B (Figure S6) imager strands consistent to inefficient triplet state depopulation of these dyes by COT^30, 32^.

**Figure 6.**
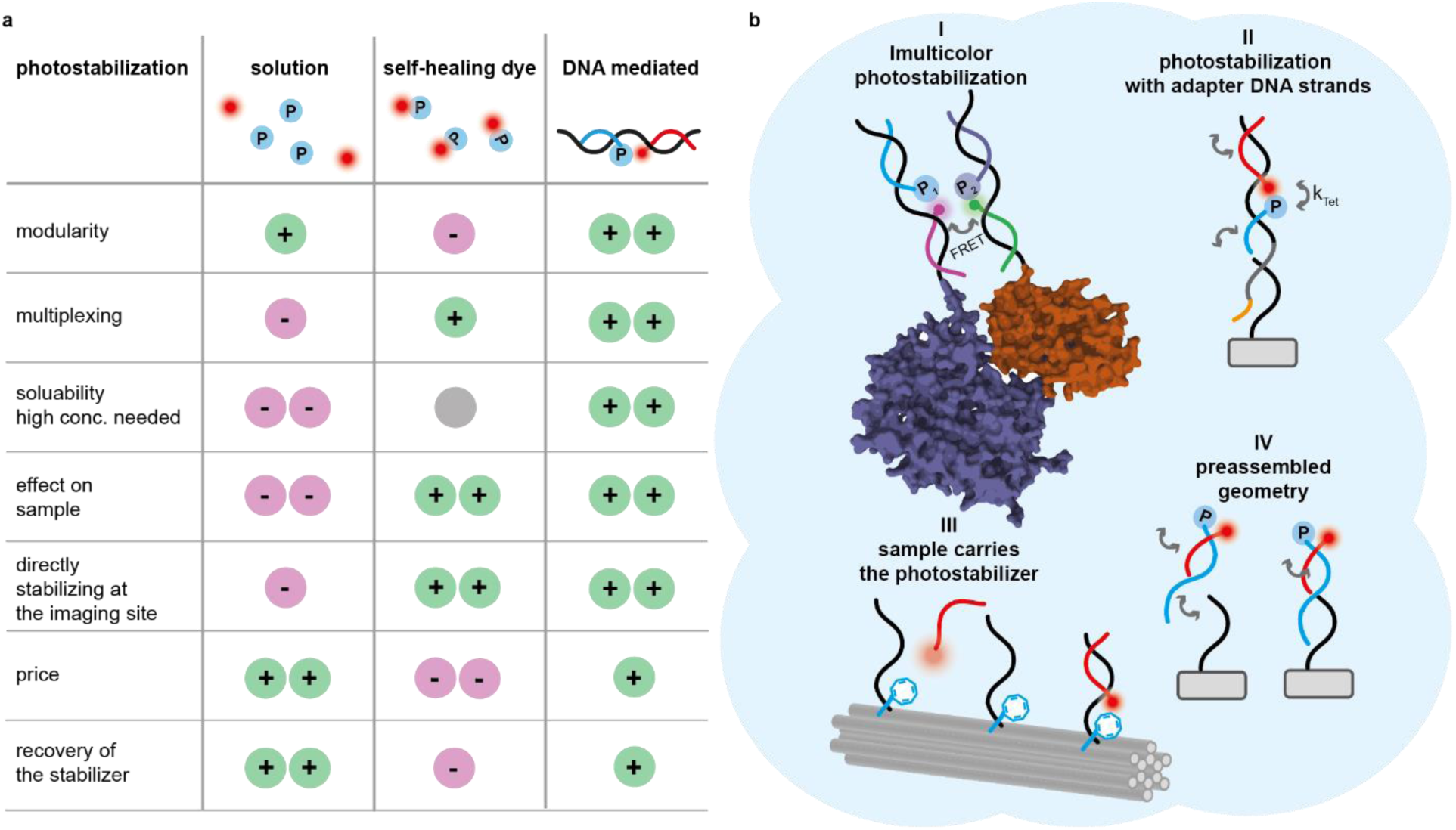
Comparison of conventional solution based photostabilization, self-healing dyes and DNA mediated photostabilization and further single-molecule imaging assays. **a)** Table stating advantages and disadvantages of common photostabilization techniques compared to the DNA-mediated photostabilization approach introduced here; **b)** Schemes showing how DNA-mediated photostabilization could be implemented in different imaging approaches: I: “mix and match” use of DNA-mediated photostabilization in multicolor imaging when two dyes need different photostabilizer molecules; II: introducing the photostabilizer molecule via the adapter strand (orange: toehold for displacement, gray: complementary to ssDNA on the target, black: for imager and stabilizer binding) used in multiplexed DNA-PAINT measurements^63, 64^; III: functionalization of biomolecule itself with the photostabilizer to place it directly at the imaging site (e.g. via DNA origami staple strand); IV: preassembly of photostabilizer and imager strands to create modular self-healing constructs.

The sequence specificity and modularity of our DNA mediated design enables multicolor or multiplex imaging while circumventing undesired cross-interactions (Figure 6a, second row). Especially when using multiple fluorophores simultaneously, one is confronted with challenges in choosing the right photostabilization approach. The DNA-mediated approach, however, is not limited to one TSQ molecule type. Any photostabilizing moiety that can be coupled to DNA, or to any other site-specific binders (e.g. in peptide-PAINT^65^), can be implemented in this approach. Since the individual photostabilizer is directed only to the specific imaging site, different fluorophores can be optimally stabilized within the same imaging solution, enabling multicolor measurements optimized for all dyes in the experiment (Figure 6a, 6bI).

To further increase multiplexing capabilities for imaging several targets in complex biological samples, our DNA-mediated photostabilization approach can also easily be combined with current adapter-mediated techniques, that use transient secondary labels for imager binding^63, 64^ (Figure 6bII). It is even conceivable to incorporate the photostabilizer molecule directly into the sample itself, e.g., by functionalizing a protruding staple strand in a DNA origami (Figure 6bIII) or coupling a short photostabilizer strand to an antibody. When designed smartly, the added linkage error can be minimal^60, 62^. Since the photostabilizer acts only locally and specifically at the imaging site of the reporting fluorophore (Figure 6a, fifth row), the contrast when compared to unspecific signal is additionally enhanced given that the directed photostabilization approach does not act on non-specifically bound labels.^66^

Coupling the photostabilizer to DNA not only allows to direct it to the specific imaging site, but also brings the additional advantage of increasing the solubility of the TSQ entity (Figure 6a, third row). Many TSQs, like COT, are poorly water-soluble organic molecules, calling for pre-dissolvement in organic solvents like DMSO^16^ or methanol^24^. This often leads to precipitation of the organic TSQs in the imaging buffer at high concentrations (typically mM range of TSQ and ca. 1% organic solvent). In our DNA mediated approach, TSQs without pre-dissolvement in an organic solvent still resulted in successful photostabilization of the fluorophore (Figure 5), highlighting the advantage of exploiting DNA as the carrier scaffold of the photostabilizer molecule and potential platform to test even less water-soluble TSQs.

These findings make the DNA mediated approach an attractive tool for minimally invasive imaging in a biological context, where the addition of high concentrations of TSQs such as COT^46^ and of organic co-solvents like DMSO^67^ can influence the sample of interest (Figure 6a, fourth row). Additionally, carefully prepared cell samples often undergo multiple imaging rounds under prolonged exposure, either for the sake of higher resolution or to deduce the interplay of several components. This makes photostabilization mandatory in most cases, but not always straightforward to implement. In our DNA-mediated approach the soluble photostabilizer can be added at a fraction (10^7^ less as in Figure 5) of the concentration needed for the solution-based technique. The stabilizing entity can also bind permanently, reducing the amount of additive in solution to zero (Figure 2 and 3). While exploiting DNA interactions allows to specifically direct the photostabilizer to the imaging site (Figure 6a, fifth row), it comes at the cost of increasing the linkage error of the fluorescent label on the object of interest. Nevertheless, the achieved 3D resolution (Figure 5) indicates, that the introduced additional linkage error does not affect the achievable resolution of the applied secondary antibody labeling^60, 62^. Additionally, for imaging applications such as DNA-PAINT based kinetic referencing^68^ or quantitative DNA-PAINT (qPAINT)^69^, a slightly increased linkage error is irrelevant to the measure outcome but a high stability of the DNA docking site is a prerequisite. The specificity of DNA-smediated photostabilization, hence, makes it an attractive tool for these super-resolution imaging applications requiring a large statistic of binding events over time for a reliable quantification.

In comparison to self-healing dyes that require multiple synthesis steps or come with high associated costs when obtained commercially, coupling of the photostabilizer to DNA is relatively simple and affordable (Figure 6a, sixth row) and the same TSQ-coupled oligonucleotide can be reused in multiple imaging schemes. As it has been shown in our experiments with pCOT (and is also the case for self-healing dyes^32^), a single photostabilizing moiety is not sufficient when higher excitation intensities are needed. In such situations the performance of the self-healing dyes would be limited by the stability of the photostabilizer. In contrast, as demonstrated in Figure 4, the DNA-mediated approach can overcome this bottleneck by photostabilizer recovery (Figure 6a, seventh row). Analogous to self-healing dyes, it is, however, also conceivable to pre-assemble the photostabilizer and imager strands (Figure 6bIV) in a stable DNA duplex^25, 27, 28, 31^, that binds to the imaging site via a single-stranded overhang. As has been recently reported, using partially double-stranded DNA could also additionally help reduce non-specific binding and, therefore, undesired background in DNA-PAINT imaging applications^66^. The pre-assembled geometry could hence serve as a cost-effective approach to emulate the self-healing dye strategy.

Currently, the performance of our DNA mediated photostabilization strategy is both restricted by DNA as the mediating agent, making it susceptible to DNA degrading conditions (e.g., DNAses), and by the imperfect photostabilizer COT. The rather low energy of its triplet state (ca. 0.8 eV^70^) only allows for photostabilization of dyes with low triplet state energies. Quenching a dye’s triplet state via energy transfer, leads to the formation of the triplet excited state of COT which has a lifetime of up to 100 µs^70^ introducing a potentially reactive long-lived intermediate and reducing the duty cycle of triplet state depopulation. An improved performance, thus, requires a TSQ entity with 1) tunable triplet state energy to extend the approach to broader range of fluorescence labels; 2) a shorter triplet state lifetime to preclude the formation of long-lived intermediates and to improve the stability of TSQ itself. We are currently exploring both avenues to create a library of DNA-mediated photostabilizers that could be applied to broad range of fluorescence labels in multicolor imaging applications.

## Conclusion and Outlook

In conclusion, we have developed a modular DNA-mediated photostabilization approach that relies on delivery of photostabilizers directly to the imaging site. We demonstrated that the approach allows to improve photon budgets of permanent dye labels at lower excitation intensities outperforming solution additives which are used at several orders of magnitude larger concentrations (Figure 2). Nevertheless, at increased excitation intensities or repetitive binding of multiple fluorophores (Figure 3), the stability of the photostabilizer itself becomes a limiting factor. To address this, we introduced the recovering photostabilization scheme (rCOT), where the photostabilizer is continuously exchanged but still acts directly at the imaging site (Figure 4). rCOT significantly slowed down the loss of DNA-PAINT localizations, even under high excitation intensities and ambient oxygen conditions. Surprisingly, introduction of rCOT to DNA-docking sites also reduced association and dissociation rates of the imager strand.

We further demonstrated the applicability of our approach to complex imaging environments by imaging microtubules in cells (Figure 5). pCOT photostabilization improved the localization rate and precision of super-resolution images, even under oxygen-rich conditions. When combined with the SIMPLER algorithm, we achieved axial resolution and 3D reconstruction capabilities comparable to those obtained with more stable and brighter dyes expanding the palette of fluorescence labels that are suited for super-resolution imaging.

Our minimally invasive photostabilization strategy offers a promising solution for challenging imaging environments where the delivery of high concentrations of additives is prohibited. The modularity of our approach enables its adaptation to various imaging schemes, facilitating the development of multicolor imaging techniques, screening of new photostabilizers, and expansion to different fluorescence labels.

## Methods

### General materials

For folding, purification and storage of 12HB DNA origami nanostructures, a 1× TAE buffer with 16 mM MgCl_2_ was used. Bleaching of permanent fluorescent labels and DNA-PAINT with DNA origami were performed in a 2× PBS buffer with 75 mM MgCl_2_. Bleaching of recovering labels was performed in a 2× PBS buffer with 500 mM NaCl and 0.05% Tween 20. ^71, 72^

Oxygen-free single-molecule imaging was performed by addition of 1% (wt/v) *D*-(+)-glucose (Sigma Aldrich, USA), 165 units/mL glucose oxidase (G2133, Sigma Aldrich, USA), and 2170 units/mL catalase (C3155, Sigma Aldrich, USA) to the imaging solution.^24^

The p8064 scaffold strand was extracted from M13mp18 bacteriophages. Unmodified staple strands were purchased from Eurofins Genomics GmbH and Integrated DNA Technology Inc. Dye labeled oligonucleotides for DNA-PAINT imaging or permanent labeling were purchased from Eurofins Genomics GmbH (Germany).

The activated COT-maleimide linker molecule was synthesized as reported previously.^52^ Labeling and purification of the COT-modified oligonucleotides was performed at Ella Biotech GmbH (Germany).

### DNA origami folding

All investigated 12HB DNA origami nanostructures (Figure S1) were folded in a 1× TAE buffer containing 16 mM MgCl_2_ using the corresponding p8064 scaffold strand extracted from M13mp18 bacteriophages with a non-linear thermal annealing ramp over 16 hours (Table S1).^73^ Concentrations of scaffold strand, unmodified and modified staple strands in the folding mix are given in Table S2. Modifications of the DNA Origami were designed using caDNAno (version 2.2.0). A full list of the unmodified staple strands and sequences of the 12HB DNA origami^74^ is given in Table S8. Folded DNA origami nanostructures were purified with 100 kDa MWCO Amicon Ultra filters (Merck, Germany). Concentrations of purified sample solutions were measured via UV/vis spectroscopy (NanoDrop, Fischer Scientific, USA). Correct folding of the origami structures was confirmed via AFM imaging (Figure S1) on a NanoWizard® 3 ultra AFM (JPK Instruments AG).

### Sample preparation

High precision 170 µm thick microscope cover glass slides (22×22 mm, Carl Roth GmbH, Germany) were initially ultrasonicated in a 1% Hellmanex solution. After thoroughly washing with ultra-pure water, the glass slides were irradiated for 30 min in a UV ozone cleaner (PSD-UV4, Novascan Technologies, USA). Cleaned glass slides and microscope slides were assembled into an inverted flow chamber as described previously.^71^ The assembled chambers were rinsed with 1× PBS, and passivated with 50 µL of BSA-biotin (0.5 mg/mL in PBS, Sigma Aldrich, USA) for 15 minutes and washed with 50 µL 1× PBS. The passivated surfaces were incubated with 50 µL Neutravidin (0.25 mg/mL in 1× PBS, Sigma Aldrich, USA) or 50 µL Streptavidin (0.5 mg/mL in 1× PBS, Sigma Aldrich, USA) for 15 minutes and washed with 50 µL 1× PBS. The sample solution with DNA origami featuring four staple strands with biotin modifications on the base (Table S8) was diluted to approximately 50 pM in 1× PBS buffer containing 500 mM NaCl and incubated in the chambers for ca. 5 minutes and stored in a 1× TAE containing 10 mM MgCl_2_. Sufficient surface density was probed with a TIRF microscope.

### Imager and photostabilizer strands

To ensure specific hybridization of the imager (labelled with Cy5) and photostabilizer (labelled with COT) oligonucleotides on a single DNA docking site, their corresponding strands were designed to have orthogonal DNA sequences. Fluorescent label strands, so-called imager strands, were designed of varying lengths (8, 11 and 20 nt) to probe permanent and recovering labeling (sequences in Table S3). The photostabilizer strands labeled with a COT moiety on the 5’-end were also of varying lengths (rCOT with 10 nt, pCOT with 17 nt) to compare permanent and recovering photostabilization (sequences in Table S4). For reference measurements without a COT moiety on the DNA docking site, a 17 nt strand with the pCOT sequence without a COT label was used. DNA docking sites, consisting of a combination of complementary sequences of one of the COT strands and one of the imager strands, were modified to the 3’-ends of selected staple strands on the 12HB nanostructure (for sequences, see Table S5-7). For more details, see SI section 1.3.

### Labeling of DNA docking sites on 12HB origami

Permanent labels, i.e., label strand with lengths of 17 or more nucleotides, were hybridized to immobilized DNA origami nanostructures by incubation of a 10 nM label solution in a 2× PBS buffer with 75 mM MgCl_2_ for 60 minutes. After washing away with 2× PBS with 500 mM NaCl and 0.05% Tween 20, the labeled DNA origami was stored in a 1× TAE containing 10 mM MgCl_2_. Recovering and shorter COT and imager oligonucleotides were added to the imaging solution at different concentrations specified in the manuscript.

### Single-molecule fluorescence imaging

Automated bleaching experiments of permanent and recovering fluorescent labels and DNA-PAINT measurements of DNA origami nanostructures were performed on a commercial Nanoimager S (ONI Ltd., UK) with red excitation at 638-nm and green excitation at 532 nm, respectively. The microscope was set to TIRF illumination and widefield movies were acquired with frames of 100 ms (bleaching of permanent labels with 0.3 or 2.0 kW/cm^2^) or 200 ms exposure time (bleaching of permanent and recovering labels with 0.1 kW/cm^2^). For more details on bleaching experiments of permanent and recovering fluorescent labels, see SI sections 1.6 and 1.7. For more details on DNA-PAINT imaging on DNA origami nanorulers, see SI section 1.8.

### Data analysis of single-molecule fluorescence trajectories

Bleaching movies were first background corrected with ImageJ 1.52n (version 1.8.0_172). Individual spots were picked and corresponding single-molecule trajectories were extracted with a custom written ImageJ script. Bleaching trajectories were then analyzed with a custom written Python script using Hidden Markov Modeling (HMM). For every bleaching curve, brightness, i.e., photon count per frame, the total number of photons before bleaching and the time point of bleaching were extracted. Apparent photon numbers were converted in absolute photon numbers using the specifications of the used sCMOS camera.

### Fluorescence correlation spectroscopy

Autocorrelation FCS studies were performed on immobilized 12HB origami nanostrucurres labeled with a permanent COT oligonucleotide (pCOT) and a permanent imager strand containing Cy5B in the presence of oxygen scavenging system. Single-molecule fluorescence trajectories of surface immobilized emitters were acquired on a home-built confocal microscope equipped with time-correlated single photon counting (TCSPC) capabilities (as described previously^75^) upon excitation with 639-nm laser (2 µW excitation intensity, measured at the objective). Single photon counting data was read into Python using a home-written script and analysed using the *pcorrelate* function of the module pycorrelate^76^. The corresponding analysis script can be found on GitLab. The function uses an algorithm described in literature^77^ to calculate the cross-correlation function between two channels at time lag τ via:

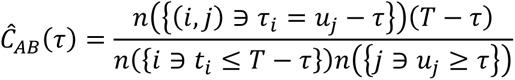

where t _i_ is the arrival time of the i^th^ photon in channel A, u_j_ is the arrival time of the j^th^ photon in channel B, n is the operator for counting the elements in the list and T is the experimental time. We calculated *Ĉ_AB_* for the timestamps collected in one channel, (“auto-correlation”, i.e. channel A = channel B). For each experimental condition, we acquired at least 19 single molecule trajectories for no COT and 43 for pCOT, calculated *Ĉ_AB_* for each trace and averaged the result.

### DNA-PAINT measurements of DNA-origami nanostructures

DNA-PAINT measurements on immobilized DNA origamis were also performed on the commercial Nanoimager S (ONI Ltd., UK). The microscope was set to TIRF illumination and an excitation power density of ca. 1.0 kW/cm^2^ at 638 nm. Widefield movies totaling 36000 frames were acquired at 100 ms time binning over 60 min.

TIFF files were analyzed using the Picasso software package.^71^ For fitting the centroid position information of single point spread functions (PSF) of individual imager strands, the MLE (Maximum Likelihood Estimation) analysis was used with a minimal net gradient of 2500 and a box size of 5. The fitted localizations were further analyzed with the “Render” module from Picasso. X-y-drift correction of the localizations was performed using RCC drift correction algorithm. Individual docking sites on the 12HB nanostructures were picked using Picasso’s “Pick tool”, setting the pick diameter to 0.6 camera pixels to extract the corresponding binding kinetics and photon statistics per docking site. To obtain accurate brightness values, the localizations of every picked docking site were filtered in order to remove the contributions from the first and the last frames of a binding event, using a custom written Python code as described previously.^58^

### Cell culture

COS-7 cells (ATCC) were cultured in DMEM (Gibco, No. 11965084) medium supplemented with 10% FBS (Gibco, No. 10500064). Cells were passaged twice a week using 0.05% trypsin EDTA (Gibco, No. 25300054).

### Preparation for microtubule imaging

COS-7 cells were seeded on Ibidi eight-well glass-bottom chambers (No. 80827) at a density of 25 000 cm^−2^. In preparation for imaging, cells were fixed using the protocol described by Whelan and Bell^78^, using 0.4% Glutaraldehyde (Sigma Aldrich, USA) and 0.25% Triton X-100 (Sigma Aldrich, USA) in CSB (1M NaCl, 100 mM PIPES, 30 mM MgCl_2_, 10 mM EGTA, 10 mM Sucrose; pH = 6.2) for 90s. After rinsing with 37°C PBS twice, 3 % Glutaraldehyde in CSB were incubated for 15 min, followed by washing with PBS (30s, 1min, 5min, 10min, 15min). The reductant NaBH_4_ was added at 0.5% (w/v) to quench residual aldehyde, followed by PBS washing steps (30s, 1min, 5min, 10min, 15min). For blocking, the cells were incubated in antibody incubation buffer (Massive Photonics) for 45 minutes. Primary rat anti-tubulin antibody (Massive Photonics) was added 1:100 and incubated overnight, after washing twice with washing buffer (WB, MP) secondary anti-rat Ab (MP) was added at 1:100 and incubated overnight, washed three times, and then stored in washing buffer. Prior to imaging, for pCOT samples, the 17 nt COT strand was incubated at 200 pM for 1.5 h at 37°C to ensure hybridization in WB + 50 mM MgCl_2_. Directly before imaging, the imager was added to the solution at 200 pM. For rCOT samples, analogously to DNA-origami measurements, the tenfold concentration (2 nM) was added, together with the 200 pM imager before the measurement.

### DNA-PAINT in fixed cells

DNA-PAINT measurements in fixed cells were carried out on a custom-built total internal reflection fluorescence (TIRF) microscope, based on an inverted microscope (IX71, Olympus) equipped with a nosepiece (IX2-NPS, Olympus) for drift suppression. For red excitation a 150 mW laser (iBeam smart, Toptica Photonics) spectrally filtered with a clean-up filter (Brightline HC 650/13, Semrock) was used. A diffractive beam shaper (piShaper 6_6_VIS, AdlOptica) generated a flat-top laser beam profile, which guaranteed a homogeneous illumination of the sample across the whole detection plane. For more details, see SI section 1.5. COS-7 samples, without and with pCOT/rCOT were measured using an excitation power density of ca. 0.6 kW/cm^2^ for 36000 frames at 100 ms exposure time and EM gain set to 150.

### Analysis of DNA-PAINT in fixed cells

From raw data photon counts and x/y coordinates were extracted using the “Localize” feature of the software Picasso. Therein, PSF fitting was performed using MLE with minimal net gradient 12000 and box size 5. To correct for drift, RCC was applied in Picasso “Render”. The drift-corrected data was subjected to filtering using a custom written software.^58^ With this, first and last frame were excluded to factor out photon count errors due to incompletely acquired binding events. Only localizations that were detected for more than three frames within half a camera pixel size (93 nm size, distance threshold 50 nm) were included in the filtered data. Exemplary rendered images were extracted at the same zoom and contrast settings for all samples and applying the individual localization precision blur. Rendered images with 32 color coding according to photon were extracted setting the maximum photon number to 10000.

To obtain 3D cross sections of microtubules, localizations were picked using the rectangular tool in Render perpendicular to the microtubules’ length. Subsequently, the previously reported custom-built SIMPLER software in MATLAB was used to extract the axial positions.^58^ We used the following parameters: N_0_ (photons expected for z = 0) = 7000, θ_i_ (incident angle) = 66°, α (evanescent component) = 0.9, NA = 1.45, λ_0_ (excitation wavelength) = 644 nm and λ_d_ (mean detection wavelength) = 700 nm. From this, the ThunderStorm plugin for Image-J was used to create the z-color coded image rendering, as reported in the SIMPLER publication, using a pixel size of 3.5 nm in the super-resolved image, where every localization is rendered as a Gaussian blurred spot with a width of 7 nm. Localization precision was calculated with a custom software, analysing individual ON-events. Here, also a minimum ON-time of 3 frames is required before calculating standard deviation in x/y and average number of photons from an event.

## Acknowledgement

V.G. thankfully acknowledges German Research Foundation (DFG, grant number GL 1079/1-1, project number 503042693); L.Z. and V.G. acknowledge Center for Nanoscience (CeNS) for a collaborative research grant; P.T. gratefully acknowledges financial support from DFG (INST 86/2224) and from the Federal Ministry of Education and Research (BMBF) and the Free State of Bavaria under the Excellence Strategy of the Federal Government and the Länder through the ONE MUNICH Project ‘Munich Multiscale Biofabrication’ and the BMBF in the framework of the Cluster4Future program (Cluster for Nucleic Acid Therapeutics Munich, CNATM) (Project ID: 03ZU1201AA). S.S.M. and M.S. were supported by the Intramural Research Program of the National Institutes of Health (NIH), NCI CCR. A.Z. acknowledges the support from Alexander von Humboldt Foundation. L.Z., T.C. and A.H. thankfully acknowledge German Research Foundation (DFG, project number 518284393).

## Author contributions

V.G. and M.S. conceived the idea. M.S. and V.G. designed and the experiments on DNA origami nanostructures. C.C., M.S., and V.G. designed DNA-PAINT experiments in fixed COS-7 cells. M.S. synthesized and characterized DNA origami nanostructures. M.S. performed bleaching experiments and DNA-PAINT imaging on DNA nanostructures and subsequent data analysis. C.C. performed DNA-PAINT imaging in fixed COS-7 cells and subsequent data analysis with help of A.M.S. M.S. and V.G. performed single-molecule fluorescence correlation studies. A.M.S. provided custom-written software and support for filtering and analysis of DNA-PAINT microtubule data. J. B. built the flat-top TIRF setup to enable DNA-PAINT cell measurements. F.C. provided custom-written software for the analysis of single-molecule bleaching data. L.G. provided custom-written software for fluorescence auto correlation analysis. A.M.S. and J.B. provided custom written software for filtering and analysis of obtained DNA-PAINT data. S.S.M. and M.J.S. synthesized and provided Cy5B-NHS ester. J.K., A.H., L.Z., and T.C. synthesized and provided the COT-maleimide linker molecule. V.G., A.M.S., and P.T. supervised the study. M.S. and C.C. visualized the data. V.G., M.S., C.C., and A.M.S. wrote the manuscript with additional input from P.T., J.B., and L.G. All authors reviewed and approved the manuscript.

