## Supporting Information for "Minimally Invasive DNA-Mediated Photostabilization for Extended Single-Molecule and Super-resolution Imaging"

‡ These authors contributed equally

### 1. Methods and Materials

#### 1.1. General materials

For folding, purification and storage of 12HB DNA origami nanostructures, a 1×TAE buffer with 16 mM $MgCl_2$ was used. Bleaching of permanent fluorescent labels and DNA-PAINT with DNA origami were performed in a 2× PBS buffer with 75 mM $MgCl_2$. Bleaching of recovering labels was performed in a 2× PBS buffer with 500 mM NaCl and 0.05% Tween 20.[1, 2]

Oxygen-free single-molecule was performed by addition of 1% (wt/v) *D*-(+)-glucose (Sigma Aldrich, USA), 165 units/mL glucose oxidase (G2133, Sigma Aldrich, USA), and 2170 units/mL catalase (C3155, Sigma Aldrich, USA) to the imaging solution.[3]

The p8064 scaffold strand for the folding of the DNA Origami nanostructures were extracted from M13mp18 bacteriophages. Unmodified staple strands were purchased from Eurofins Genomics GmbH and Integrated Device Technology Inc. Dye labeled oligonucleotides for DNA-PAINT imaging or permanent labeling were purchased from Eurofins Genomics GmbH (Germany).

The COT-maleimide compound was synthesized by the Cordes Group as previously reported.[4] Labeling of the COT-maleimide to thiol modified DNA was performed at Ella Biotech GmbH (Germany).

Specific materials used for individual experiments are described in the sections below.

#### 1.2. DNA Origami folding

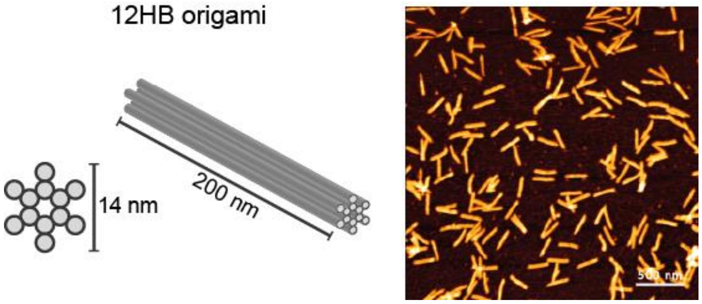

**Figure S1.** Scheme of the 12HB DNA origami used in this study and exemplary AFM scan of purified 12HB illustrating the successful self-assembly of the designed structures.

**Table S1**. Thermal ramp used for the folding of the 12HB origami nanostructures.

| Temperature (°C) | Time per °C (min) | Temperature (°C) | Time per °C (min) |
|---|---|---|---|
| 65 | 2 | 44 | 75 |
| 64 – 61 | 3 | 43 | 60 |
| 60 – 59 | 15 | 42 | 45 |
| 58 | 30 | 41-39 | 30 |
| 57 | 45 | 38-37 | 15 |
| 56 | 60 | 36-30 | 8 |
| 55 | 75 | 29-25 | 2 |
| 54-45 | 90 | 4 | storage |

**Table S2.** Final concentrations and relative equivalents of scaffold strand, unmodified staple strands (core staple strands) and modified staple strands (e.g. biotinylated staple strands for immobilization and DNA-PAINT docking site staple strands for superresolution imaging) used within this study.

| Reagent | Final concentration [nM] | Equivalents |
|---|---|---|
| Scaffold strand | 20 | 1 |
| Core staple strands | 200 | 10 |
| Docking site staple strands | 600 | 30 |
| Biotinylated staple strands | 600 | 30 |

#### 1.3. Imager and Photostabilizer strands

All used imager strand sequences are given in Table S3. Three different imager strands all labelled with a fluorophore on the 3'-end, have been employed within this study. To investigate permanent fluorescent labels, a 20 nt long imager strand was used to label to a hybridize to a 20 nt docking site sequence. To investigate the photostability of a recovering label, a 11 nt imager sequence was used as reported previously.[5] For DNA-PAINT imaging, a 8 nt subsequence of the 20 nt permanent sequence was used.

For permanent labeling, the green fluorophores Cy3 and Cy3B and the red fluorophores Cy5 and Atto647N were labelled to the pImg strand. For further investigation of a recovering label, the red fluorophore Cy5 was labelled to the rLabel strand. For DNA-PAINT imaging, the red fluorophores Cy5 and Cy5B were labelled to the 8 nt long fImg strand.

**Table S3.** Fluorescently labelled imager strands used within this study. A 20 nt long permanent imager (pImg) and 11 nt long recovering label (rLabel) were used for bleaching experiments of permanent and recovering labels. DNA-PAINT imaging was performed with a 8 nt long fast imager strand (fImg). All imager strands were labelled with fluorophores on their 3'-end.

| Name | Length (nt) | Sequence (5` to 3`) |
|---|---|---|
| pImg | 20 | TATGAGAAGTTAGGAATGTT-Dye |
| fImg | 8 | GGAATGTT-Dye |
| rLabel | 11 | TTTCCCTTTTT-Dye |

All used COT DNA strands are given in Table S4. To investigate permanent and dynamic COT strands, the COT-maleimide compound was coupled to the 5'-end of a thiolated DNA oligonucleotide (Scheme **S1**) with varying sequence lengths (17 nt for permanent pCOT strand and 10 nt for recovering rCOT strand).

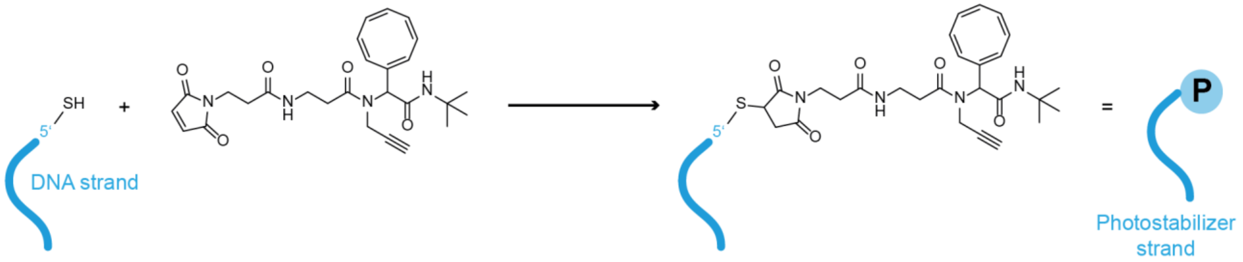

**Scheme S1.** Coupling of maleimide functionalized COT linker molecule to thiolated DNA oligonucleotide resulting in a photostabilizer strand with the COT entity at the 5' end.

**Table S4.** COT labelled photostabilizer strands used in this study. For a permanent label, COT was labelled to a 17 nt long permanent strand (pCOT). For a dynamic labeling, COT was modified to a 10 nt fast exchanging photostabilizer strand (rCOT). All photostabilizer strands were labelled with COT on their 5'-end.

| Name | Length (nt) | Sequence (5` to 3`) |
|---|---|---|
| pCOT | 17 | COT-ATGATGTAGGTGGTAGA |
| rCOT | 10 | COT-ATGATGTAGG |

#### 1.4. Widefield TIRF microscopy

Automated bleaching experiments of permanent and recovering fluorescent labels and DNA-PAINT on DNA origami nanostructures were performed on a commercial Nanoimager S (ONI Ltd., UK). Red excitation at 638 nm was realized with a 1100 mW laser, green excitation at 532 nm with a 1000 mW laser, respectively. The microscope was set to TIRF illumination. In order to not corrupt the first frames of the acquired intensity transients by the photobleaching of single DNA origami nanostructures, the objective was first focused into the sample plane on a random section of the glass surface and the auto focus was activated. Subsequently the imaging lasers were shut off. Before starting time lapse measurements, the sample slide was moved to a new region of interest while still being kept in focus by the auto focus. The data acquisition was initialized by activating the lasers and taking frames of 100 ms to 200 ms over a user defined acquisition protocol.

DNA-PAINT measurements in fixed cells were carried out on a custom-built total internal reflection fluorescence (TIRF) microscope, based on an inverted microscope (IX71, Olympus) equipped with a nosepiece (IX2-NPS, Olympus) for drift suppression. For yellow excitation, a 560 nm/1 W fiber laser (MPB Communications) filtered with a clean-up filter (Brightline HC 561/4, Semrock) was used. Red excitation at 644 nm was realized with a 150 mW laser (iBeam smart, Toptica Photonics) spectrally filtered with a clean-up filter (Brightline HC 650/13, Semrock). The red and the yellow beams were coupled into polarization maintaining single mode fibers (P3-488PM-FC-2 for 560 nm, P3-630PM-FC-2 for 644 nm) to obtain perfect Gaussian beam profiles. Behind the fibers, the excitation beam paths were combined with a dichroic mirror (T612lpxr, Chroma). To obtain a homogenous excitation profile across the whole detection plane, the laser light was guided through a diffractive beam shaper (piShaper 6_6_VIS, AdlOptica) that changes the Gaussian beam profile to a flat-top beam profile. The laser beam was coupled into the microscope body with a triple-color beam splitter (Chroma z476-488/568/647, AHF Analysentechnik) and focused on the back focal plane of an oil-immersion objective (100×, NA = 1.45, UPlanXApo, Olympus) with a telescope, that could be aligned for TIRF illumination. An additional ×1.6 optical magnification lens was applied to the detection path resulting in an effective pixel size of 92.6 nm. The fluorescence light was spectrally cleaned up (ET 700/75, Chroma for red excitation or ET 605/70m, Chroma for yellow excitation) and recorded by an electron multiplying charge-coupled device camera (Ixon X3 DU-897, Andor), which was controlled with the software Micro-Manager 1.4.[6, 7]

#### 1.5. Surface-Immobilization of DNA origami nanorulers

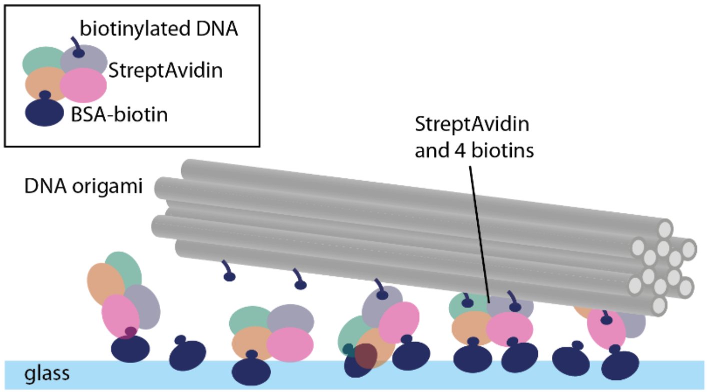

**Figure S2.** Scheme of components involved in surface immobilization of DNA origami.

#### 1.6. DNA mediated photostabilization of a permanent single-molecule label

To study the photostabilization of a permanent single-molecule label by DNA mediated collision with a COT bound to the same DNA docking site, a staple strand in the central region of the 12HB was modified at the 3'-end (Table S5) with the complementary sequences of the permanent COT strand and permanent imager strand given in Table S3 and Table S4.

After immobilization of DNA origami on neutravidin functionalized glass slides, 10 nM of the COT strand and 10 nM of the permanent imager strand were incubated in a 2×PBS with 500 mM NaCl and 0.05% w/w Tween® 20 for 60 min and excessive label strands were washed away afterwards. For bleaching experiments, the photostabilization buffer was applied to the sample chambers. Bleaching was performed under low (0.3 kW/cm$^2$) and high (2.0 kW/cm$^2$) excitation power to investigate photostability in different excitation regimes. For low excitation powers, 3000 frames of 200 ms were acquired over an overall observation period of 10 min. For high excitation powers, 600 frames of 100 ms were acquired over an overall observation period of 1 min.

DNA mediated photostabilization was probed for two permanent green (Cy3, Cy3B) and two permanent red fluorophore labels (Atto647N, Cy5).

**Table S5**. Modified staple strand in the central region of the 12HB for DNA mediated photostabilization of permanent fluorescent labels Sequences are denoted from 5'- to 3'-end. The docking site staple strand exhibits a 17 nt binding sequence for the pCOT strand, marked in blue, and a 20 nt binding sequence for a permanent imager strand, marked in red, respectively. The numbers for the 5'- end 3'-end of the staples represent the helix number in the corresponding caDNAno file. Number in brackets represent the starting and ending position of the staple in the corresponding helix.

| Name | Docking Length (nt) | Site | Sequence (5` to 3`) | 5'-end ´ | 3'-end] |
|---|---|---|---|---|---|
| pCOT + pImg | 17 + 20 | | TCGTTCACCGCCTGGCCCT-TCTACCACCTACATCAT-AACATTCCTAACTTCTCATA | 10[331] | 11[344] |

#### 1.7. DNA mediated photostabilization of a recovering single-molecule label

To study the photostabilization of a recovering single-molecule label by DNA mediated collision with a COT bound to the same DNA docking site, a staple strand in the central region of the 12HB was modified at the 3'-end (Table S6) with the complementary sequences of the permanent or dynamic COT strand (10 or 17 nt) and recovering imager strand (11 nt) given in Table S3 and Table S4.

Permanent COT strand was labelled to DNA origami immobilized on streptavidin functionalized glass slides by incubation of a 10 nM pCOT strand solution in a 2×PBS with 500 mM NaCl and 0.05% w/w Tween® 20 for 60 min. Dynamic rLabel strands labelled with Cy5 (10 nM) and fast exchanging rCOT strands (100 nM) were added to the photostabilizing imaging buffer with 500 mM NaCl and 0.05% w/w Tween®.

Photostability of the recovering label and the DNA docking site was probed under low excitation power (0.1 kW/cm$^2$) over 18000 frames of 200 ms over an overall observation period of 60 min.

**Table S6.** Modified staple strand in the central region of the 12HB for DNA mediated photostabilization of recovering fluorescent labels Sequences are denoted from 5'- to 3'-end. The docking site staple strand exhibits a 10 or 17 nt binding sequence for the rCOT or pCOT strand, marked in blue, and a 11 nt binding sequence for a recovering imager strand, marked in red, respectively. The numbers for the 5'- end 3'-end of the staples represent the helix number in the corresponding caDNAno file. Number in brackets represent the starting and ending position of the staple in the corresponding helix.

| **Name** | **Docking Site Length (nt)** | **Sequence (5` to 3`)** | **5'-end ´** | **3'-end]** |
|---|---|---|---|---|
| pCOT + rLabel | 17 + 11 | TCGTTCACCGCCTGGCCCT-TCTACCACCTACATCAT-AAAAAGGGAAA | 10[331] | 11[344] |
| rCOT + rLabel | 10 + 11 | TCGTTCACCGCCTGGCCCT-CCTACATCAT-AAAAAGGGAAA | 10[331] | 11[344] |

#### 1.8. DNA-PAINT imaging on DNA origami nanorulers

To study the applicability of the DNA mediated photostabilization for super-resolution microscopy, three staple strands with ca. 90 nm distances on the 12HB were modified at the 3'-end (Table S7).with the complementary sequences of the permanent or dynamic COT strand (10 or 17 nt) and fast imager strand (8 nt) given in Table S3 and Table S4.

Permanent COT strand was labelled to DNA origami immobilized on streptavidin functionalized glass slides by incubation of a 10 nM pCOT strand solution in a 2×PBS with 500 mM NaCl and 0.05% w/w Tween® 20 for 60 min. The 8 nt fast imager strand (1 nM) and fast exchanging rCOT strand (100 nM) were added to the photostabilizing imaging buffer with 75 mM $MgCl_2$.

Photostability of the DNA-PAINT docking sites was probed under high excitation power (1.2 kW/cm$^2$) over 36000 frames of 100 ms over an overall observation period of 60 min.

**Table S7.** Modified staple strands with 90 nm distances on the 12HB for DNA-PAINT imaging using DNA mediated photostabilization. Sequences are denoted from 5'- to 3'-end. The docking site staple strands exhibit a 10 or 17 nt binding sequence for the rCOT or pCOT strand, marked in blue, and an 8 nt binding sequence for the fast imager strand, marked in red, respectively. The numbers for the 5'- end 3'-end of the staples represent the helix number in the corresponding caDNAno file. Number in brackets represent the starting and ending position of the staple in the corresponding helix.

| **Name** | **Docking Site Length (nt)** | **Sequence (5` to 3`)** | **5'-end ´** | **3'-end]** |
|---|---|---|---|---|
| pCOT + fImg | 17 + 8 | GTATGTGAAATTGTTATCC-TCTACCACCTACATCAT-AACATTCC | 10[79] | 11[92] |
| | | TACCTGGTTTGCCCCAGCA-TCTACCACCTACATCAT-AACATTCC | 10[373] | 11[386] |
| | | AACACCCTAAAGGGAGCCC-TCTACCACCTACATCAT-AACATTCC | 10[625] | 11[638] |
| fCOT + fImg | 10 + 8 | GTATGTGAAATTGTTATCC-CCTACATCAT-AACATTCC | 10[79] | 11[92] |
| | | TACCTGGTTTGCCCCAGCA-CCTACATCAT-AACATTCC | 10[373] | 11[386] |
| | | AACACCCTAAAGGGAGCCC-CCTACATCAT-AACATTCC | 10[625] | 11[638] |

#### 2. Supporting Figures

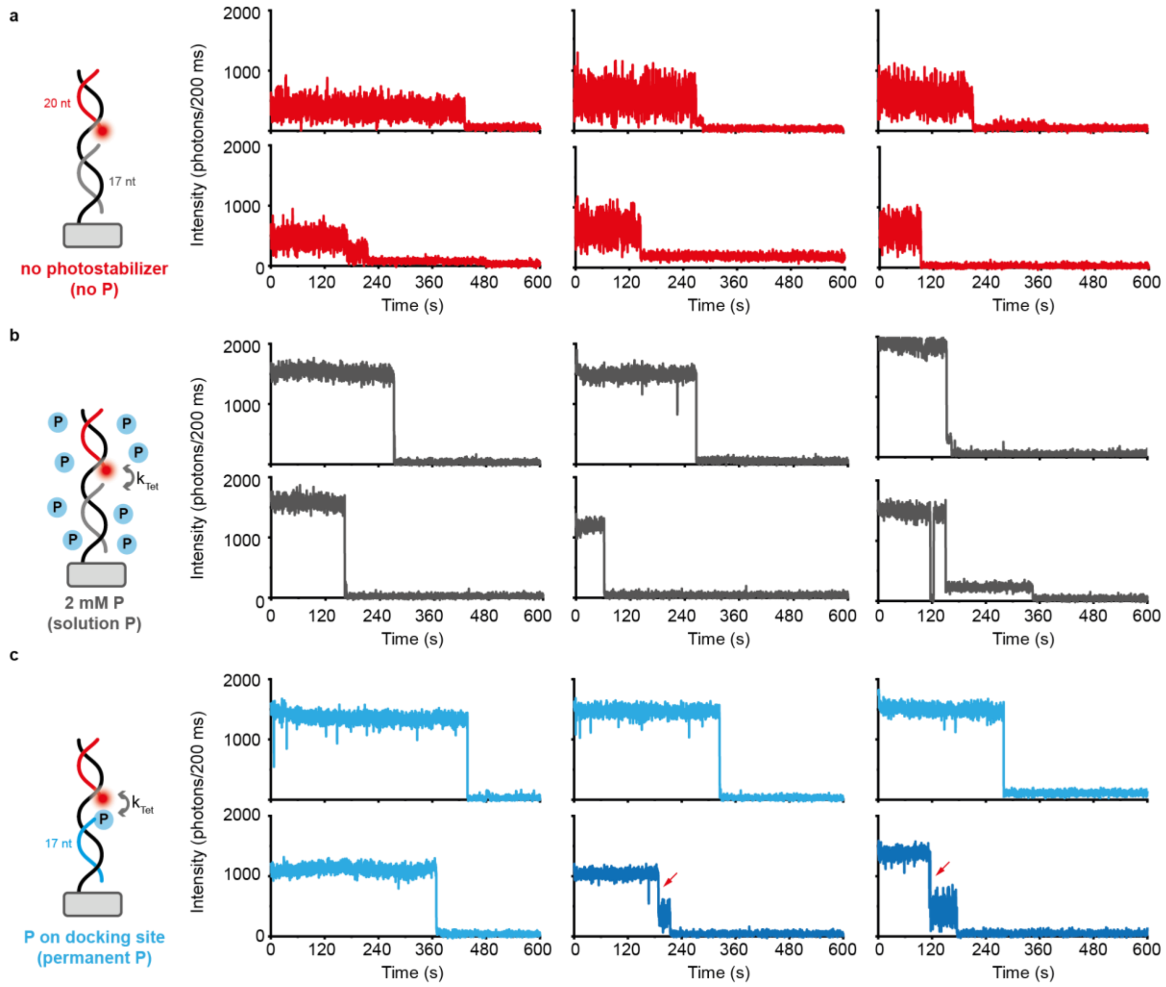

**Figure S3.** Permanent Cy5 labels with and without COT on docking site under low excitation power (0.3 kW/cm$^2$). **a)** Scheme and exemplary single-molecule trajectories for individual label spots of Cy5 labels without COT on docking site. **b)** Scheme and exemplary single-molecule trajectories for individual label spots of Cy5 labels with 2 mM COT in solution. **c)** Scheme and exemplary single-molecule trajectories for individual labels spots of Cy5 with a permanent COT label on the docking site.

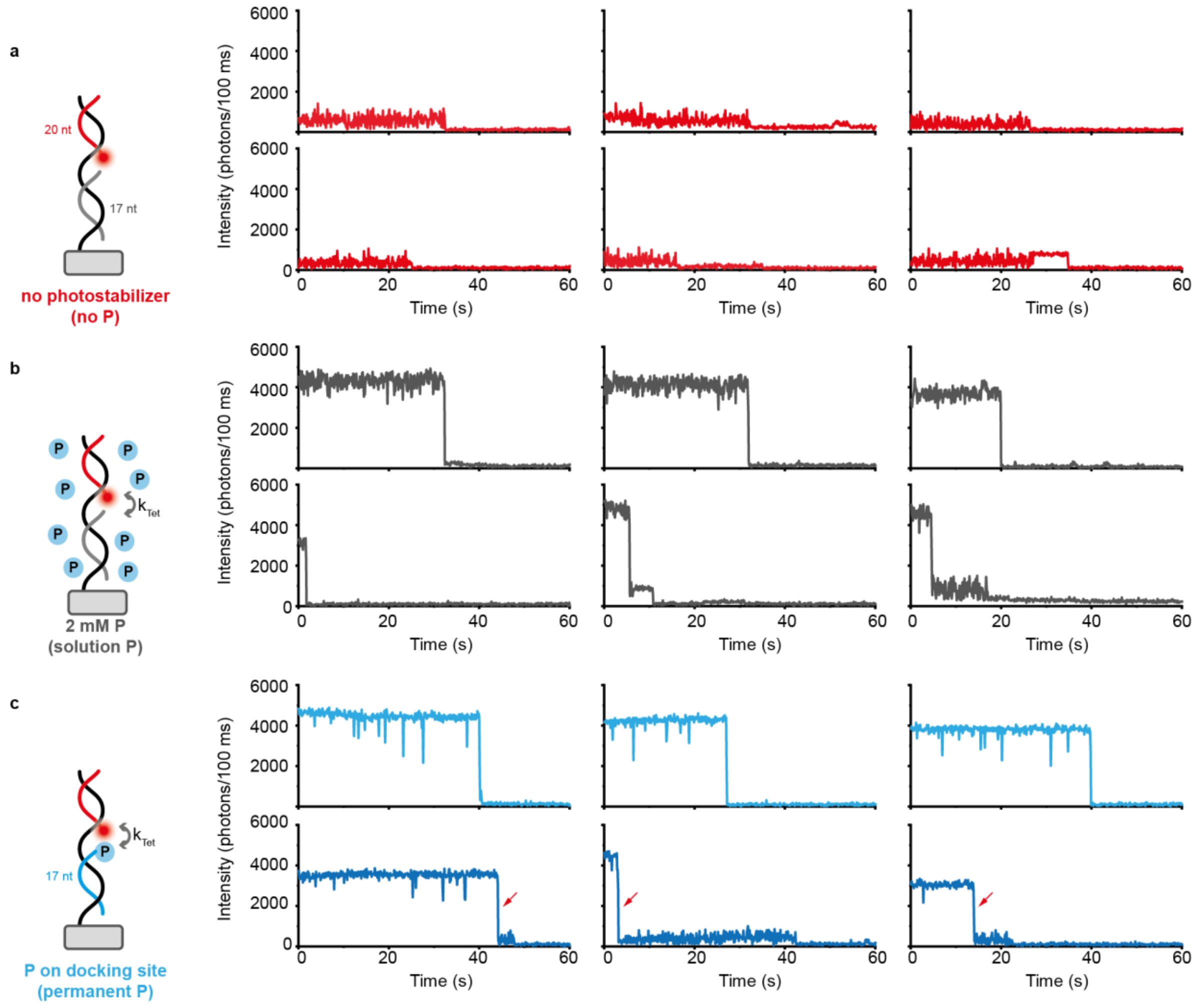

**Figure S4**. Permanent Cy5 labels with and without COT on docking site under high excitation power (2.0 kW/cm$^2$). **a)** Scheme and exemplary single-molecule trajectories for individual label spots of Cy5 labels without COT on docking site. **b)** Scheme and exemplary single-molecule trajectories for individual label spots of Cy5 labels with 2 mM COT in solution. **c)** Scheme and exemplary single-molecule trajectories for individual labels spots of Cy5 with a permanent COT label on the docking site.

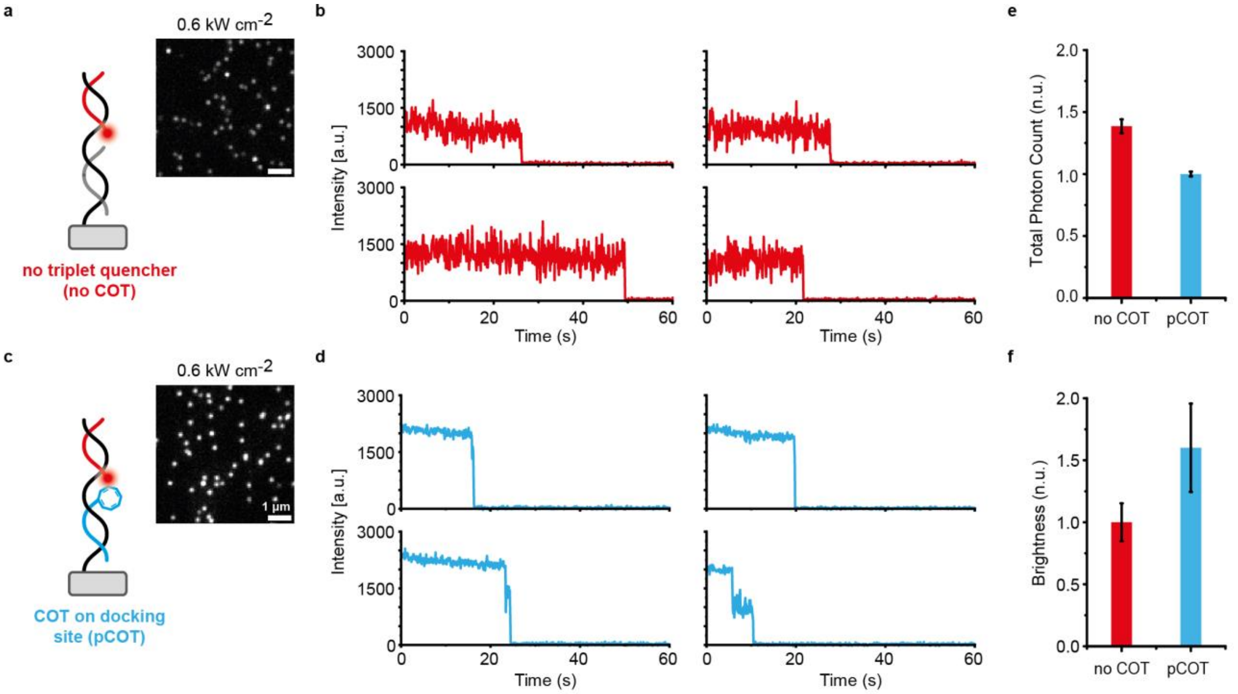

**Figure S5.** Permanent Cy3 labels with and without COT on docking site under medium excitation power (0.6 kW/cm$^2$). **a)** Scheme and exemplary TIRF image of Cy3 labels without COT on docking site. **b)** Exemplary single-molecule trajectories for individual label spots of Cy3 labels without COT on docking site. **c)** Scheme and exemplary TIRF image of Cy3 labels with a permanent COT label on the docking site. **d)** Exemplary single-molecule trajectories for individual labels spots of Cy3 with a permanent COT label on the docking site. **e)** Normalized total photon counts for permanent Cy3 labels with and without COT label on the docking site. Error bars represent error of the fit. **f)** Normalized brightness for permanent Cy3B labels with and without COT label on the docking site. Error bars represent standard deviation of gaussian fit.

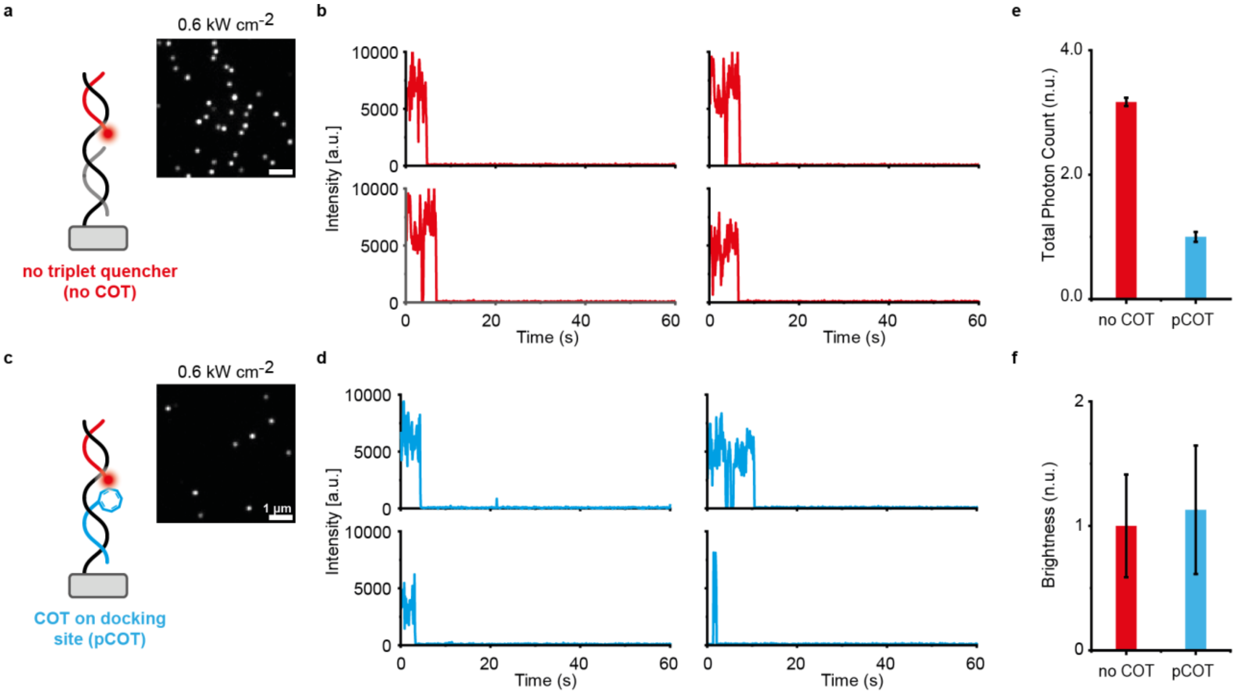

**Figure S6.** Permanent Cy3B labels with and without COT on docking site under medium excitation power (0.6 kW/cm$^2$). **a)** Scheme and exemplary TIRF image of Cy3B labels without COT on docking site. **b)** Exemplary single-molecule trajectories for individual label spots of Cy3B labels without COT on docking site. **c)** Scheme and exemplary TIRF image of Cy3B labels with a permanent COT label on the docking site. **d)** Exemplary single-molecule trajectories for individual labels spots of Cy3B with a permanent COT label on the docking site. **e)** Normalized total photon counts for permanent Cy3B labels with and without COT label on the docking site. Error bars represent error of the fit. **f)** Normalized brightness for permanent Cy3B labels with and without COT label on the docking site. Error bars represent standard deviation of gaussian fit.

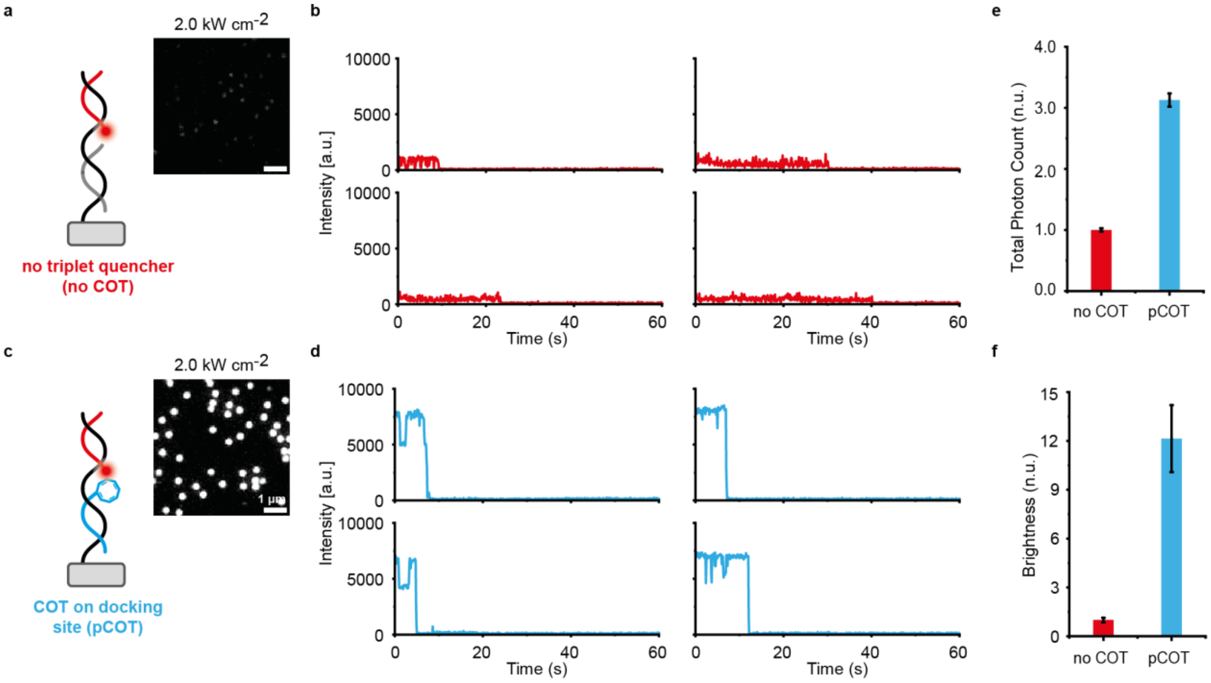

**Figure S7**. Permanent Atto647N labels with and without COT on docking site under high excitation power (2.0 kW/cm$^2$). **a)** Scheme and exemplary TIRF image of Atto647N labels without COT on docking site. **b)** Exemplary single-molecule trajectories for individual label spots of Atto647N labels without COT on docking site. **c)** Scheme and exemplary TIRF image of Atto647N labels with a permanent COT label on the docking site. **d)** Exemplary single-molecule trajectories for individual labels spots of Atto647N with a permanent COT label on the docking site. **e)** Normalized total photon counts for permanent Atto647N labels with and without COT label on the docking site. Error bars represent error of the fit. **f)** Normalized brightness for permanent Atto647N labels with and without COT label on the docking site. Error bars represent standard deviation of gaussian fit.

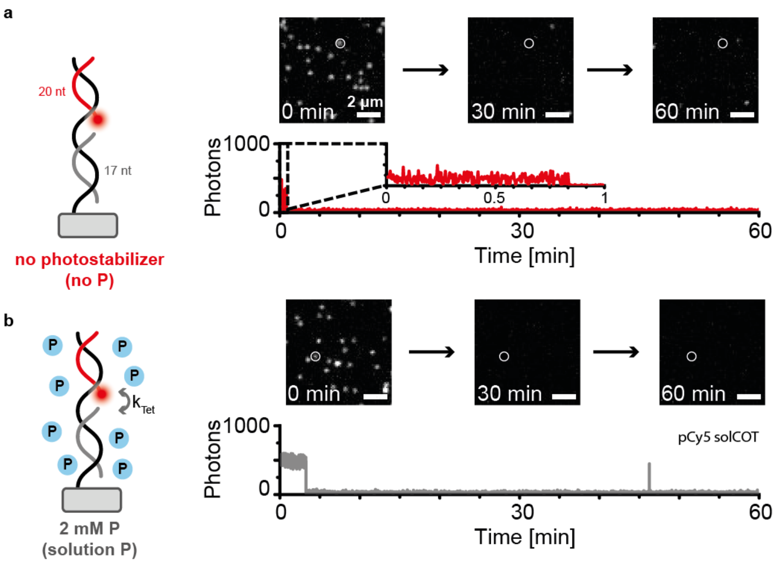

**Figure S8**. Permanent Cy5 labels without triplet state quencher (no COT) and stabilized by 2 mM COT solution under low excitation power (0.1 kW/cm$^2$). **a)** Permanent Cy5 label without triplet state quencher imaged over 60 min. **b)** Permanent Cy5 label with 2 mM COT in solution imaged over 60 min.

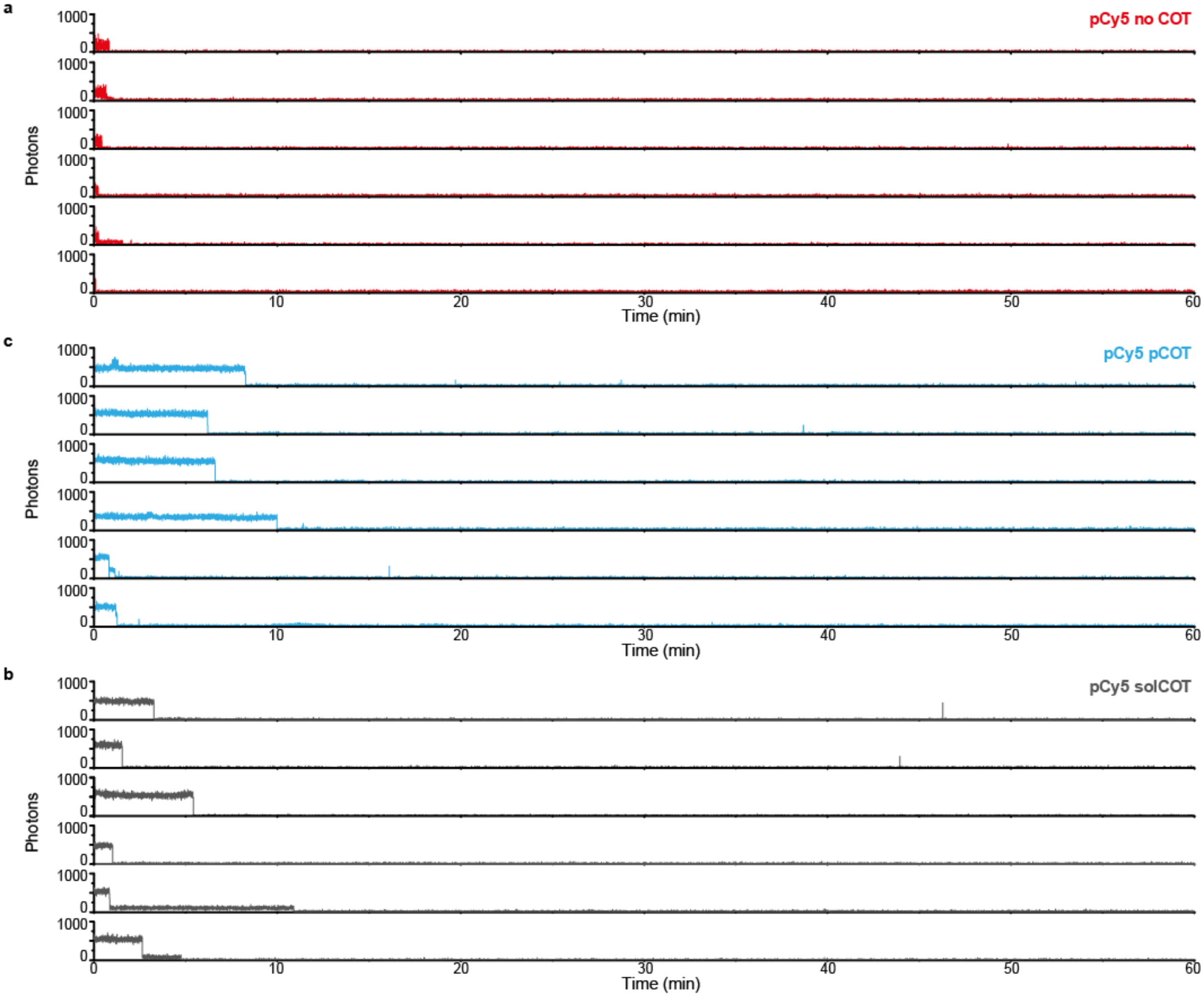

**Figure S9.** Exemplary single-molecule trajectories of permanent Cy5 labels with and without COT label on the docking site under low excitation power (0.1 kW/cm$^2$). over 60 min. **a)** Exemplary single-molecule trajectories for individual labels spots of Cy5 without COT label on the docking site (pCy5 no COT). **b)** Exemplary single-molecule trajectories for individual labels spots of Cy5 with a permanent COT label on the docking site (pCy5 pCOT). **c)** Exemplary single-molecule trajectories for individual labels spots of Cy5 with 2 mM COT in solution (pCy5 solCOT).

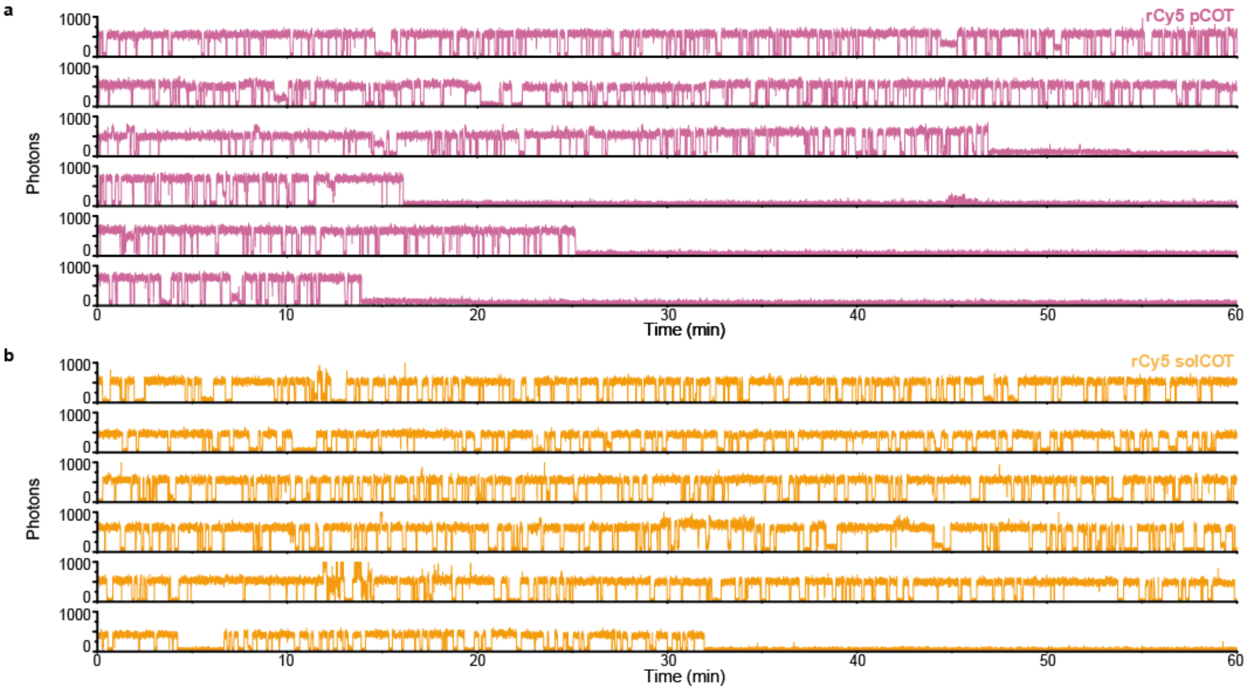

**Figure S10**. Exemplary single-molecule trajectories of recovering Cy5 labels with COT label on the docking site or in solution under low excitation power (0.1 kW/cm$^2$). over 60 min. **a)** Exemplary single-molecule trajectories for individual labels spots of recovering Cy5 with a permanent COT label on the docking site (rCy5 pCOT). **b)** Exemplary single-molecule trajectories for individual labels spots of recovering Cy5 with 2 mM COT in solution (rCy5 solCOT).

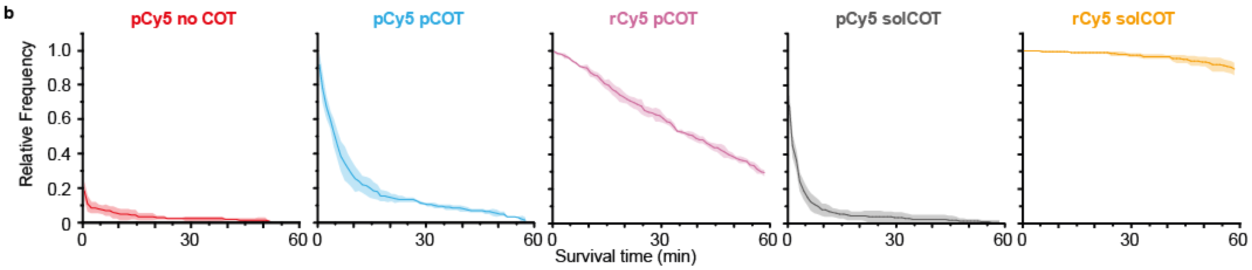

**Figure S11.** Survival times of permanent and recovering Cy5 labels with and without COT label on the docking site under low excitation power (0.1 kW/cm$^2$). Lines represent average of XXX measurements, areas represent the standard deviation.

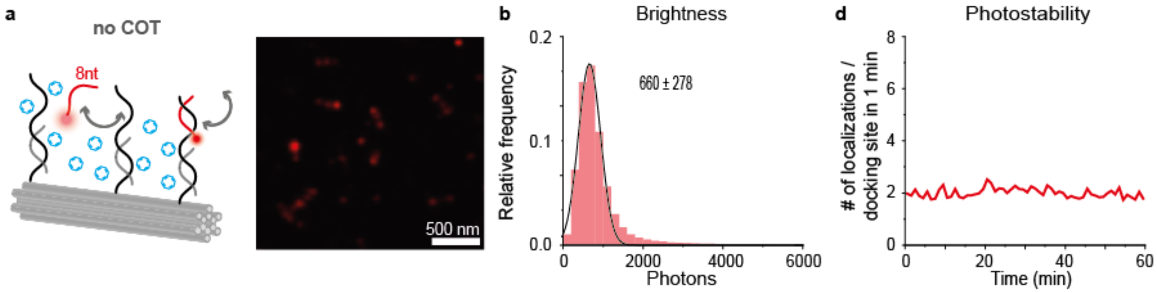

**Figure S12.** DNA-PAINT pick statistics with Cy5 imager and no COT on the docking site under high illumination power (1.0 kW/cm$^2$). **a)** Scheme of DNA-PAINT without COT (no COT) and obtained DNA-PAINT image after 60 min. **b)** Obtained photon counts for DNA-PAINT with Cy5 without COT. **c)** DNA-PAINT kinetics, i.e. on- and off-times, for individual DNA-PAINT docking sites without COT. **d)** Observed photostability of DNA-PAINT docking sites over 60 min without COT.

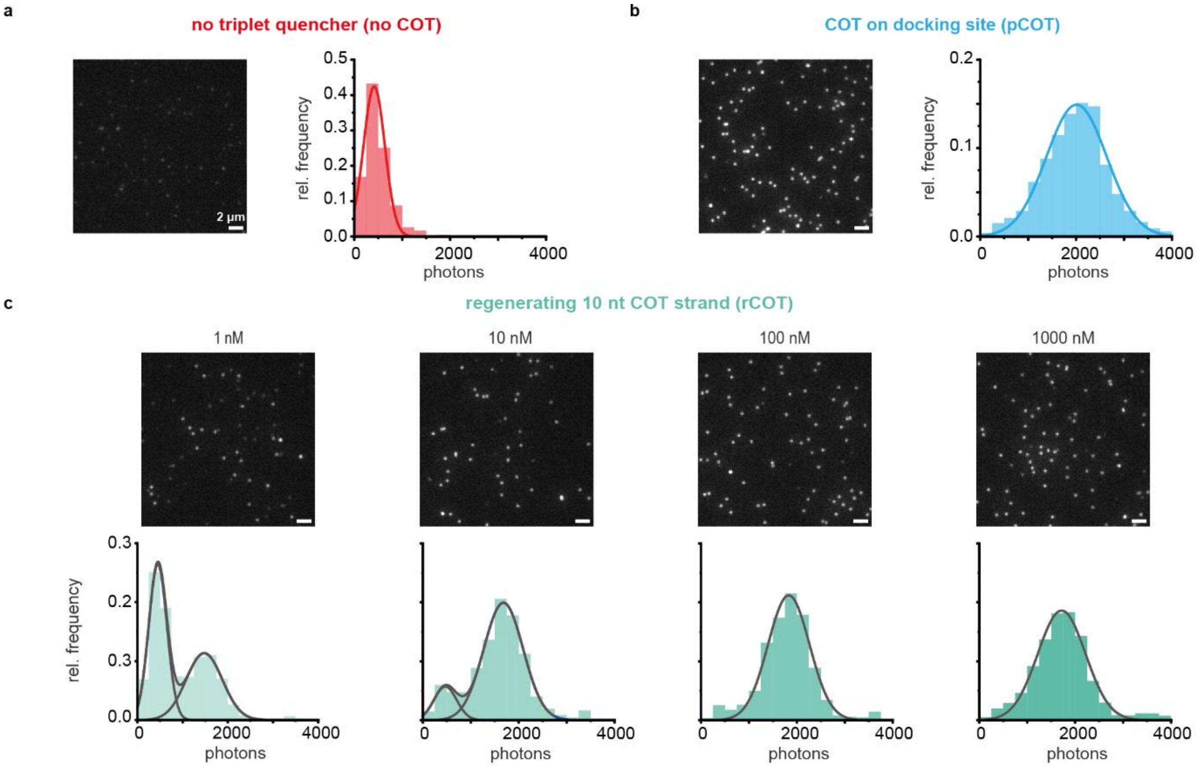

**Figure S13.** Saturating the DNA docking site with a recovering COT strand. **a)** Exemplary TIRF image of permanent Cy5 label and no COT on the docking site and extracted brightness histogram. **b)** Exemplary TIRF image of permanent Cy5 label and a permanent COT label on the docking site and extracted brightness histogram. **c)** Exemplary TIRF images of permanent Cy5 label and varying concentrations (1 nM – 1000 nM) of a recovering COT label (10 nt) on the docking site and extracted brightness histograms. All data were acquired under medium high excitation power (0.8 kW/cm$^2$).

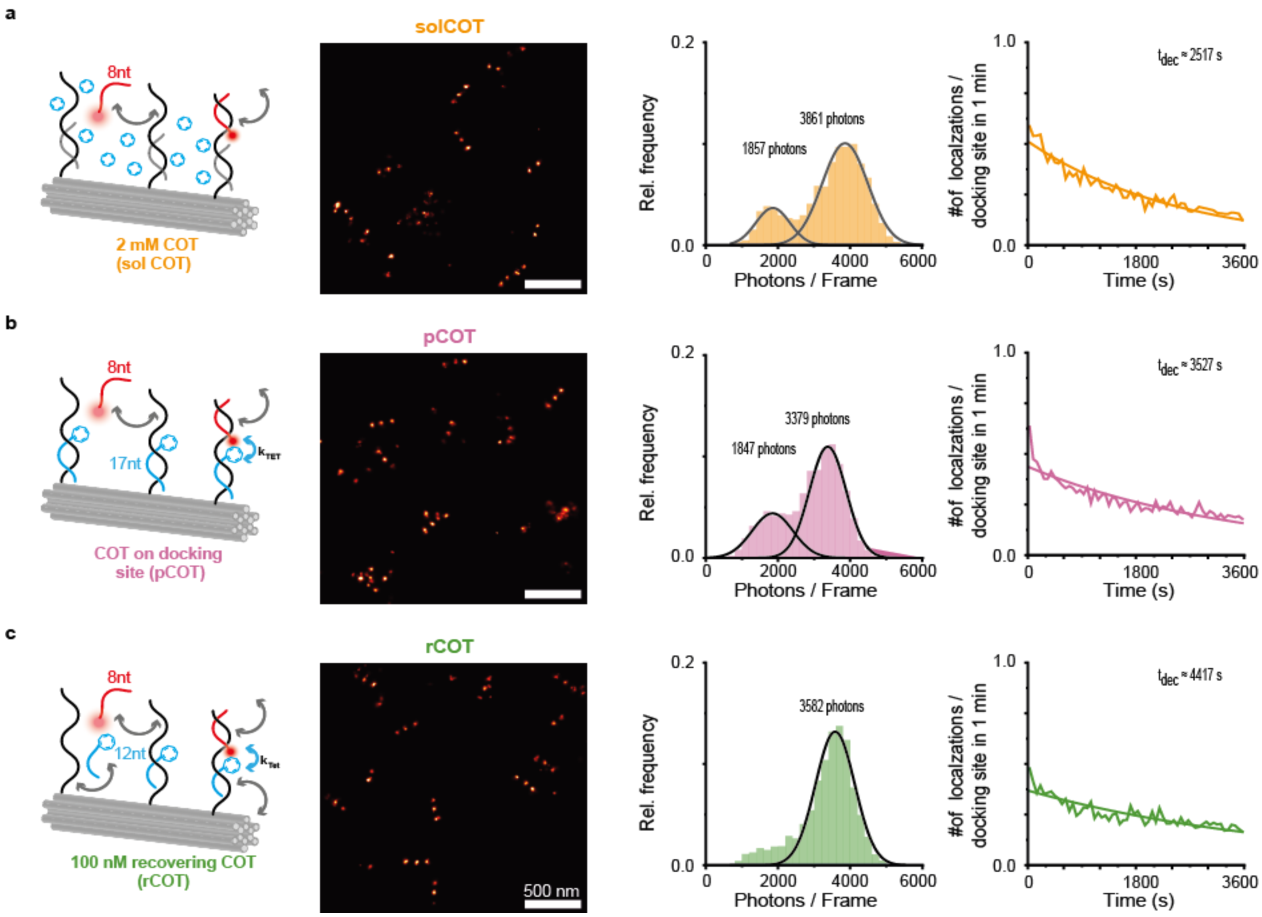

**Figure S14**. DNA-PAINT with solution-based photostabilization or DNA mediated photostabilization using COT and Cy5 under high illumination power (1.0 kW/cm$^2$) in the presence of oxygen. Exemplary reconstructed DNA-PAINT images of 3x1 12HB nanorulers, brightness values of picked DNA docking sites and detected localizations per docking site over time obtained with **(a)** 2 mM COT solution-based photostabilization, **(b)** a permanent DNA mediated photostabilization (21 nt binding sequence) and **(c)** a self-regenerating DNA mediated 10 nt self-regenerating COT label, respectively.

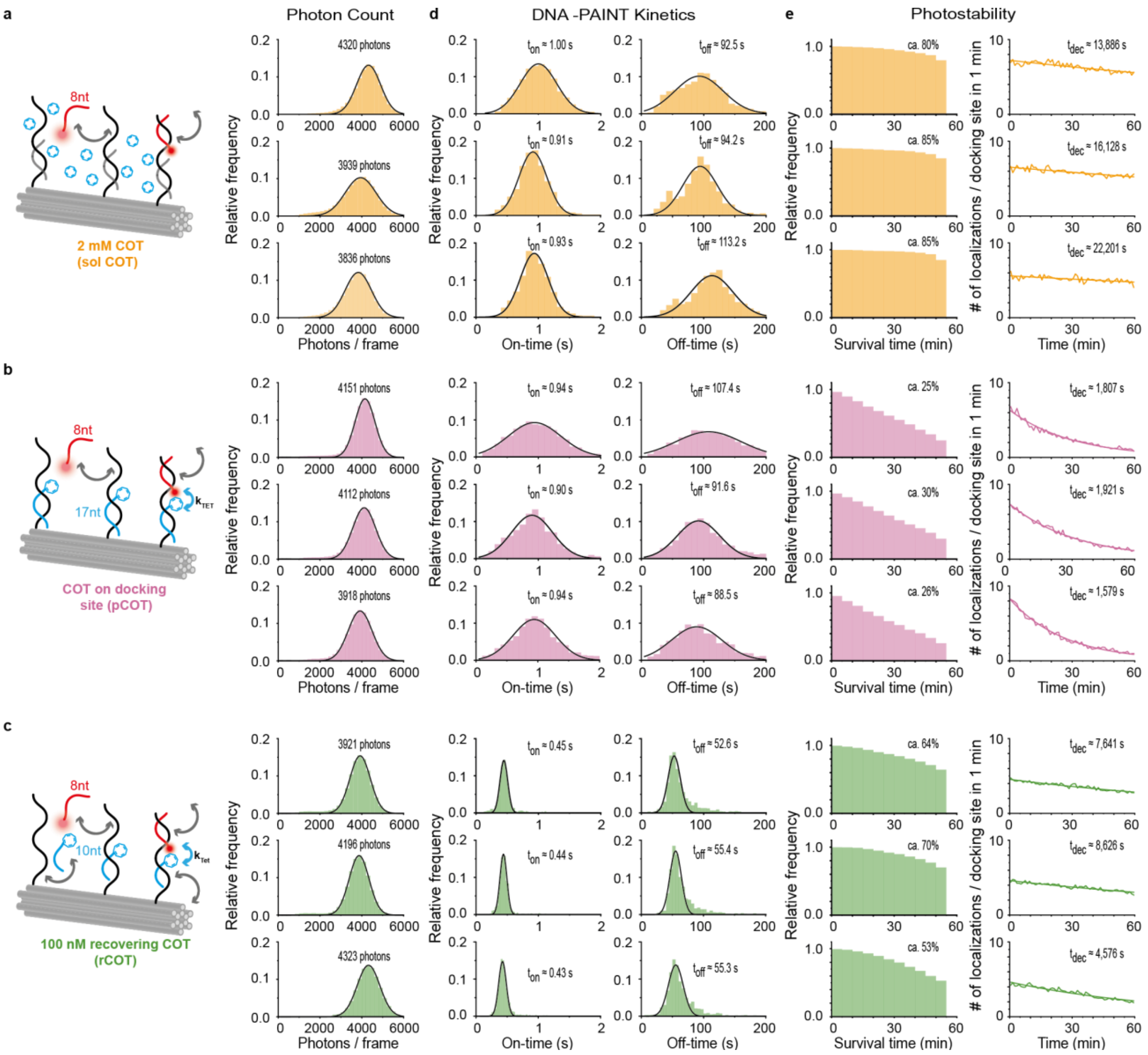

**Figure S15.** Triplicates and DNA-PAINT pick statistics with Cy5 and solution-based photostabilization vs. DNA mediated photostabilization under high illumination power (1.0 kW/cm$^2$). **a)** Scheme of DNA-PAINT with 2 mM COT in solution (solCOT) and obtained photon counts for three individual measurements. **b)** Scheme of DNA-PAINT with a permanent COT label (17 nt, pCOT) on the docking site and obtained photon counts for three individual measurements. **c)** Scheme of DNA-PAINT with a fastly recovering COT label (10 nt, rCOT) on the docking site and obtained photon counts for three individual measurements. **d)** DNA-PAINT kinetics, i.e. on- and off-times, for individual DNA-PAINT docking sites. **e)** Observed photostability of DNA-PAINT docking sites over 60 min.

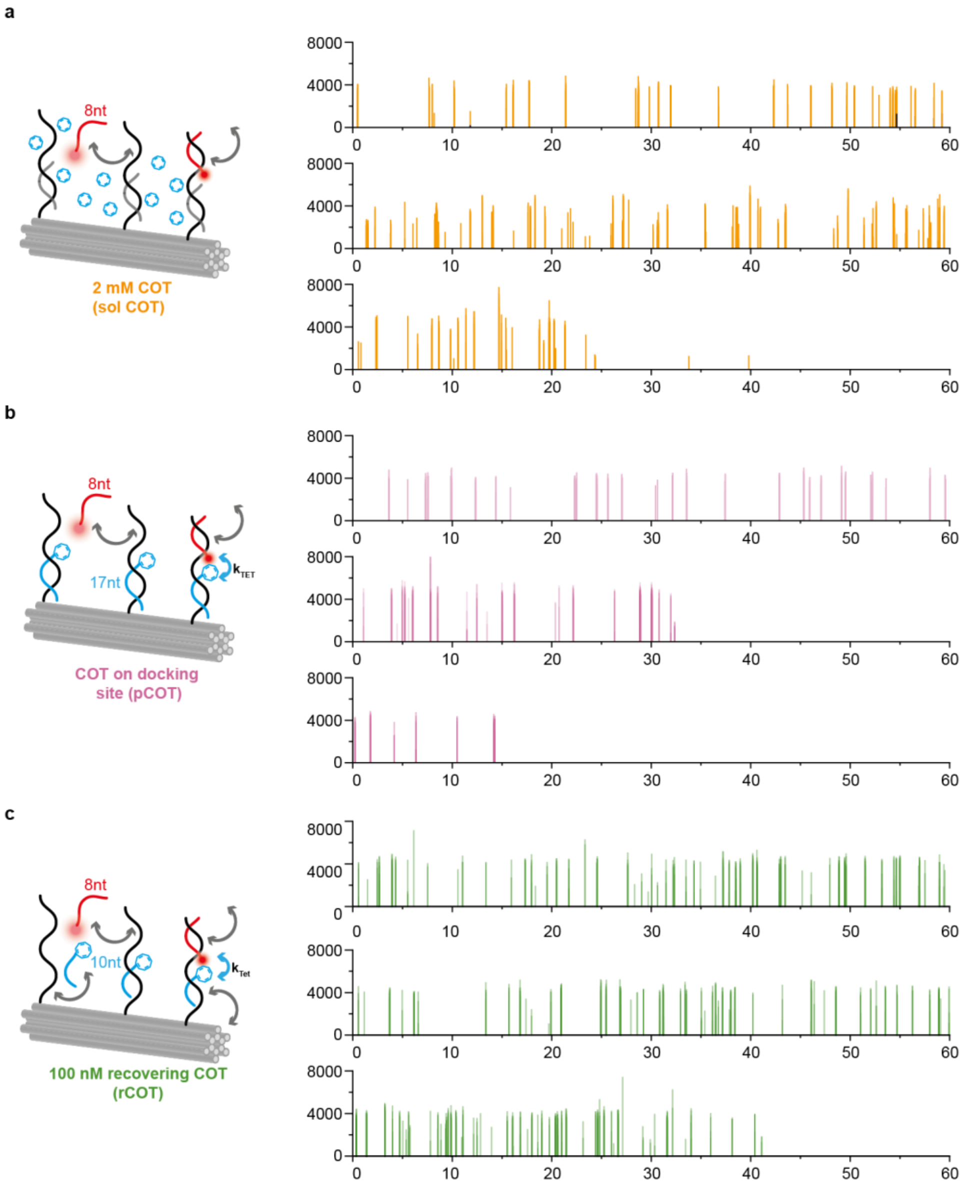

**Figure S16.** Exemplary single-molecule trajectories of individual DNA-PAINT docking sites with 2 mM COT in solution (**a**), with a permanent COT label on the docking site (**b**) and with a fastly recovering COT label on the docking site (c).

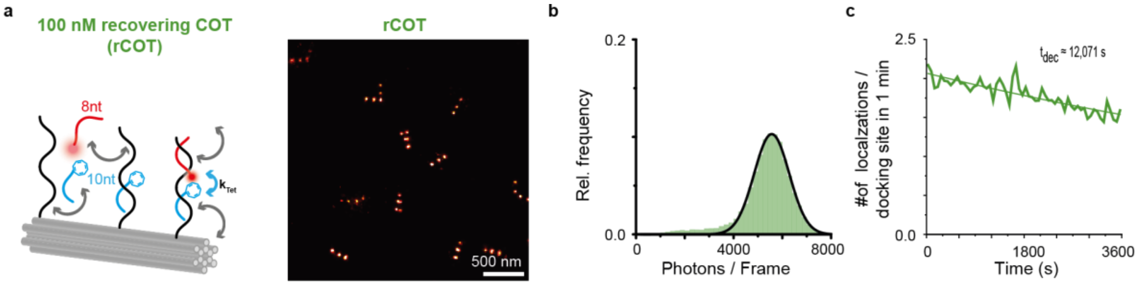

**Figure S17.** DNA-PAINT with Cy5B imager and fastly recovering COT on the docking site under high illumination power (1.0 kW/cm$^2$). **a)** Scheme of DNA-PAINT with rCOT and Cy5B and obtained DNA-PAINT image after 60 min. **b)** Obtained photon counts for DNA-PAINT with Cy5B and rCOT. **c)** Observed photostability of DNA-PAINT docking sites over 60 min with Cy5B and rCOT.

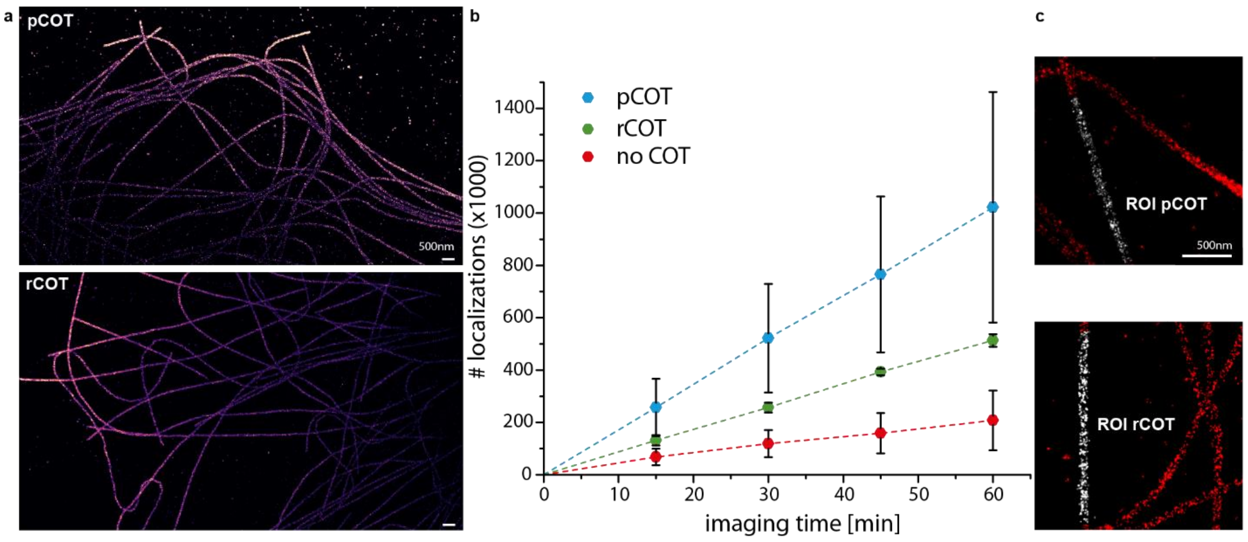

**Figure S18.** Comparison of permanent COT to recovering COT during 60 min imaging under ambient oxygen conditions. **a)** Overview zoom-in of two representative regions in the sample. **b)** Development of localizations over time for all three conditions. **c)** Exemplary ROIs from which the number of localizations was determined.

### Appendix

**Table S8.** Unmodified staple strands of 12HB DNA origami. Sequences are denoted from 5'- to 3'-end. The numbers for the 5'- end 3'-end of the staples represent the helix number in the corresponding caDNAno file. Number in brackets represent the starting and ending position of the staple in the corresponding helix.

| Unmodified staple strands | 5'-end | 3'-end |
|---|---|---|
| AAAGGGCGCTGGCAAGTATTGGC | 11[681] | 10[668] |
| GCGCCTGAATGCCAACGGCCCAGCCTCCCGCGTGCCTGTTCTTCTTTTT | 7[42] | 8[25] |
| TTGACGGGGAAAGCTTCACCAGAAATGGCATCACT | 11[651] | 6[658] |
| CATTCAACCCAAAATGTAGAACCCTCATGAATTAGTACAACC | 9[147] | 5[160] |
| TCAGAGGTGTGTCGGCCAGAATGAGTGCACTCTGTGGT | 4[60] | 7[62] |
| GGCATAAGCGTCTTCGAGGAAACGCA | 8[466] | 9[482] |
| TACATAAATTCTGGGCACTAACAACT | 8[634] | 9[650] |
| CAATCCAAAATACTGAACAGTAG | 3[457] | 10[458] |
| CATAGTTAATTTGTAAATGTCGC | 3[541] | 10[542] |
| GAACAAGAGTCCACCAATTTTTTAGTTGTCGTAGG | 11[483] | 6[490] |
| TTGAAGCCCTTTTTAAGAAAAGT | 7[441] | 7[463] |
| AAGCACAGAGCCTAATTATTGTTAGCGATTAAGACTCCTT | 7[464] | 8[448] |
| GATGTTTTTCTTTTCACCA | 10[289] | 11[302] |
| GGTCACGCCAGCACAGGAGTTAG | 3[373] | 10[374] |
| TGAACAGCTTGATACCGATAGTT | 8[363] | 8[341] |
| AAAATTCCATTCAGGCTTTTGCAAAA | 8[256] | 9[272] |
| TCCCATCCTAATGAGAATAACAT | 0[496] | 0[474] |
| ATCAGCGGGGTCAGCTTTCAGAG | 3[56] | 3[78] |
| TTCGCTATTCGCAAGACAAAGTTAATTTCATCTTC | 5[539] | 4[546] |
| TTGAGAATATCTTTCCTTATCACTCATCGAGAACA | 5[497] | 4[504] |
| GGGCGTGAAATATTAGCGCCATTCGC | 8[130] | 9[146] |
| GGCGCCCCGCCGAATCCTGAGAAGTGAGGCCGATTAAAGG | 3[667] | 0[665] |
| TTTTTTGTTTAATAAAGTAATTC | 3[476] | 3[498] |
| AAATCAGCCAGTAATAACACTATTTTTGAAGCCTTAAATC | 7[506] | 8[490] |
| AGCACTAAATCGGATCGTATTTAGACTTATATCTG | 11[609] | 6[616] |
| GGTGCCGTCGAGAGGGTTGATAT | 8[405] | 8[383] |
| GTCAGAATCAGGCAGGATTCGCG | 3[205] | 10[206] |
| TTTTTTATAACGTGCTTTCCTCTTTATAACAGTACTAT | 2[698] | 3[678] |
| AGACGGGAGAATTGACGGAAATT | 0[454] | 0[432] |
| TAAGCCAGAGAGCCAGAAGGAAACTCGATAGCCGAACAAA | 4[480] | 7[482] |
| CGCCTGACGGTAGAAAGATTCTAATGCAGATACAT | 5[245] | 4[252] |
| CAGTCTTGATTTTAAGAACTCAACGTTGCGTAT | 0[263] | 11[272] |
| CATAGAATTTGCGGTTTGAAAGAGGA | 8[298] | 9[314] |

| | | |
|---|---|---|
| GCGCAGCGACCAGCGATTATATATCATCGCCTGAT | 5[287] | 4[294] |
| TTTTTAAAAACGCTCATGGAAATA | 8[698] | 8[679] |
| AATCAGTTAAAACGTGGGAGAAA | 3[121] | 10[122] |
| AGACAACCTGAACAGTATTCGAC | 3[625] | 10[626] |
| TTTGCAACCAGCTTACGGCGGTGGTGAGGTTTCAGTTGAGGATCCTTTTT | 3[25] | 10[29] |
| TGCAACACTATCATAACCCTCGT | 7[231] | 7[253] |
| AACGAACCTCCCGACTTGCGGGA | 8[531] | 8[509] |
| CCGAACGGTGTACAGACCAGGCG | 8[321] | 8[299] |
| ATTCAAGGGGAAGGTAAATGTGGCAAATAAATC | 0[431] | 11[440] |
| GTCACCAGTACAAGGTTGAGGCA | 3[350] | 3[372] |
| TAAATCGGTTGGTGCACATCAAAAATAA | 6[153] | 2[140] |
| AGACGGCGAACGTGGCGAG | 10[667] | 11[680] |
| CCCTTCATATAAAAGAACGTAGAGCCTTAAAGGTGAATTA | 11[429] | 0[413] |
| AACTTTAATCATGGGTAGCAACG | 3[266] | 3[288] |
| ACCATCACCCAAATAAACAGTTCATTTGATTCGCC | 11[567] | 6[574] |
| TGCCTAATGAGTGAGAAAAGCTCATATGTAGCTGA | 11[147] | 6[154] |
| TTTTTTGGTAATGGGTAACCATCCCACTTTTT | 1[21] | 2[25] |
| GGAGCAGCCACCACCCTTCGCATAACGACAATGACAACAA | 7[338] | 8[322] |
| AAAAGTGTCAGCAACAATTGCAGGCGCT | 6[69] | 2[56] |
| GGTTTGCGCATTTTAACGCGAGGCGT | 8[508] | 9[524] |
| AAAAGAATAGCCCGATACATACGCAGTAAGCTATC | 11[441] | 6[448] |
| TTTCACGAGAATGACCATTTTCATTTGGTCAATAACCTGT | 7[212] | 8[196] |
| TCGGTCATACCGGGGGTTTCTGC | 8[69] | 8[47] |
| CCTCCGAAATCGGCAAAAT | 10[415] | 11[428] |
| TTCCATTGACCCAAAGAGGCTTTGAGGA | 2[307] | 3[307] |
| ACGCGTCGGCTGTAAGACGACGACAATA | 2[517] | 3[517] |
| GTCCGTCCTGCAAGATCGTCGGATTCTCTTCGCATTGGACGA | 9[105] | 5[118] |
| GTCAGTCGTTTAACGAGATGGCAATTCA | 6[615] | 2[602] |
| GAGCTTAAGAGGTCCCAATTCTGCAATTCCATATAACAGT | 4[228] | 7[230] |
| GCAGCACTTTGCTCTGAGCCGGGTCACTGTTGCCCTGCGGCTTTTT | 10[48] | 0[21] |
| TACCTGGTTTGCCCCAGCA | 10[373] | 11[386] |
| AATGCTGTAGCTGAGAAAGGCCG | 4[209] | 4[187] |
| CTATATTAAAGAACGTGGA | 10[499] | 11[512] |
| CGGTAGTACTCAATCCGCTGCTGGTCATGGTC | 0[53] | 11[62] |
| CTTGAAAACACCCTAACGGCATA | 3[247] | 10[248] |
| AAGTAAGAGCCGCCAGTACCAGGCGG | 8[382] | 9[398] |
| AAAAGATAGGGTTGAGTGT | 10[457] | 11[470] |
| TTCGCCATAAACTCTGGAGGTGTCCAGC | 2[55] | 3[55] |

| | | |
|---|---|---|
| AGGGCGAAAAACCGATTTAACGTAGGGCAAATACC | 11[525] | 6[532] |
| CCCACATGTGAGTGAATAACTGATGCTTTTAACCTCCGGC | 11[555] | 0[539] |
| TTTTTAGGAGCGGGCGCTAGGAAGGGAAGAAAGCGAATTTTT | 10[702] | 11[702] |
| TGCCATACATAAAGATTAACTGAACACCAACAGCCGGAATAG | 9[441] | 5[454] |
| TTTTTCCGGTGCAGCACCGATCCCTTACACTTGCC | 5[29] | 4[52] |
| ACAGCTGATTGCCCGTCGCTGCGCCCACACGTTGA | 11[315] | 6[322] |
| ATTAAAATAAGTGCGACGATTGGCCTTG | 2[391] | 3[391] |
| AAAACGAAAGAGGCTCATTATAC | 0[286] | 0[264] |
| TGTCCAAGTACCAGAAACCCCAG | 3[499] | 10[500] |
| TTACCAATAAGGCTTGCAGTGCGGAAGTTTAGACTGGATA | 7[254] | 8[238] |
| TTAGTGTGAATCCCTCTAATAAAACGAAAGAACGATGAATTA | 9[231] | 5[244] |
| ATCAGAGCCTTTAACGGGGTCTTAATGCCCCCTGC | 5[371] | 4[378] |
| TTACCTCTTAGCAAATTTCAACCGATTG | 6[447] | 2[434] |
| AAAACGGAATACCCAAAAGAACT | 8[489] | 8[467] |
| GTCCACGCGCCACCTCACCGTTGAAACA | 11[364] | 6[364] |
| TTTTTATCCAGCGCAGTGTCACTGC | 7[21] | 7[41] |
| GATGAATAAATCCTGTAGGTGAGGCGGTAGCGTAAGTCCTCA | 9[609] | 5[622] |
| GCTAAATCGGTTTGACTATTATA | 3[182] | 3[204] |
| CAGCTTTGAATACCAAGTTACAA | 7[567] | 7[589] |
| GGTTGCTTTGACGAGCACGTTTTT | 3[679] | 3[698] |
| CATGCCAGTGAGCGCTAATATCCAATAATAAGAGC | 5[455] | 4[462] |
| TATGCATTACAGAGGATGGTTTAATTTC | 2[265] | 3[265] |
| ACTGCCCGCTTTCCTGAAAAGCTATATTTTAAATA | 11[189] | 6[196] |
| TGATTTAGAAAACTCAAGAGTCAATAGT | 6[573] | 2[560] |
| TGGGCGCCAGGGTGATTCATTAGAGTAACCTGCTC | 11[273] | 6[280] |
| TGCAACTCAAAAGGCCGTACCAAAAACA | 6[195] | 2[182] |
| AAATAGGTAATTTACAAATAAGAAACGA | 2[475] | 3[475] |
| TGTTCCAACGCTAACGAACAAGTCAGCAGGGAAGCGCATT | 11[471] | 0[455] |
| GTGCCTGCTTTAAACAGGGAGAGAGTTTCAAAGCGAACCA | 11[219] | 0[203] |
| GTTTGATGGTGGTTCAGAACCCCGCCTCACAGAAT | 11[399] | 6[406] |
| TCACCGTCACCGGCGCAGTCTCT | 0[412] | 0[390] |
| AGACGTCGTCACCCTCAGATCTTGACGCTGGCTGACCTTC | 7[296] | 8[280] |
| TTTAGCAAACGCCACAATATAACTATATTCCCTTATAAATGG | 9[525] | 5[538] |
| AGCGTATCATTCCACAGACCCGCCACAGTTGCAGCAAGCG | 0[347] | 11[363] |
| GTATGTGAAATTGTTATCC | 10[79] | 11[92] |
| CCGAACTTTAATAAAAGCAAAGCGGATT | 2[223] | 3[223] |
| GTGAGTTAAAGGCCGCTGACACTCATGAAGGCACCAACCT | 11[303] | 0[287] |
| GCGCCCGCACCCTCTCGAGGTGAATT | 8[340] | 9[356] |

| | | |
|---|---|---|
| ACAGTTTTTCAGATTTCAATTACCGTCGCAGAGGCGAATT | 4[606] | 7[608] |
| TTTAGAACGCGAATTACTAGAAAACTATAAACACCGGAAT | 4[564] | 7[566] |
| TGACCTAAATTTTTAAACCAAGT | 4[545] | 4[523] |
| TAAAGAGGCAAAATATTTTATAA | 3[163] | 10[164] |
| GTTTACCGCGCCCAATAGCAAGC | 7[483] | 7[505] |
| TACCGGGATAGCAATGAATATAT | 3[331] | 10[332] |
| AAATTGTGTCGAGAATACCACAT | 4[293] | 4[271] |
| AAATGCGTTATACAAATTCTTAC | 8[573] | 8[551] |
| CAGATATAGGCTTGAACAGACGTTAGTAAAGCCCAAAAATTT | 9[315] | 5[328] |
| TAAGATCTGTAAATCGTTGTTAATTGTAAAGCCAACGCTC | 7[548] | 8[532] |
| CATTCTATCAGGGCGATGG | 10[541] | 11[554] |
| CTCCAATTTAGGCAGAGACAATCAATCAAGAAAAATAATA | 11[513] | 0[497] |
| GAGACAAAGATTATCAGGTCATTGACGAGAGATCTACAAA | 4[186] | 7[188] |
| AGGGACAAAATCTTCCAGCGCCAAAGAC | 2[433] | 3[433] |
| AAAATTTTTTAAAATGAGCAAAAGAA | 8[592] | 9[608] |
| CATCGGGAGAAATTCAAATATAT | 4[587] | 4[565] |
| ATCATTTACATAAAAGTATCAAAATTATAAGAAACTTCAATA | 9[567] | 5[580] |
| GCTACGACAGCAACTAAAAACCG | 3[289] | 10[290] |
| TTAGGTTGGGTTATAGATAAGTC | 0[538] | 0[516] |
| TATTGCCTTTAGCGTCAGACTGT | 7[399] | 7[421] |
| TTTTTCCGGGTACCGAGCTCGAATTCGTAATCTGGTCA | 11[29] | 10[49] |
| CTAAAGACTTTTAGGAACCCATG | 3[308] | 3[330] |
| GTGGAACGACGGGCTCTCAACTT | 3[79] | 10[80] |
| TCAGGTGAAATTTCTACGGAAACAATCG | 6[111] | 2[98] |
| AAGACGCTGAGACCAGAAGGAGC | 3[560] | 3[582] |
| AGCAGTCGGGAAACCTGTC | 10[205] | 11[218] |
| AACAACATGTTCATCCTTGAAAA | 3[518] | 3[540] |
| ATAATGAATCCTGAGATTACGAGCATGTGACAAAAACTTATT | 9[483] | 5[496] |
| GAGGTAACGTTATTAATTTTAAAACAAATAATGGAAGGGT | 11[597] | 0[581] |
| ACCGCATTCCAACGGTATTCTAAGCGAGATATAGAAGGCT | 4[522] | 7[524] |
| CAGCATCAACCGCACGGCGGGCCGTT | 8[46] | 9[62] |
| GCTCAAGTTGGGTAACGGGCGGAAAAATTTGTGAGAGATA | 11[93] | 0[77] |
| GGAATCGGAACATTGCACGTTAA | 3[583] | 10[584] |
| ATAAGAAGCCACCCAAACTTGAGCCATTATCAATACATCAGT | 9[399] | 5[412] |
| GGCGACACCACCCTCAGGTTGTACTGTACCGTTCCAGTAA | 11[387] | 0[371] |
| CATGTCAGAGATTTGATGTGAATTACCT | 6[279] | 2[266] |
| AATAGCTGTCACACGCAACGGTACGCCAGCGCTTAATGTAGTA | 9[651] | 5[664] |
| GCAGCACCGTAAGTGCCCGTATA | 4[419] | 4[397] |

| | | |
|---|---|---|
| ATGAATCCCAGTCACGATCGAACGTGCCGGCCAGAGCACA | 7[86] | 8[70] |
| TATGTGATAAATAAGGCGTTAAA | 7[525] | 7[547] |
| TTAATGAATCGGCCATTCATTCCAATACGCATAGT | 11[231] | 6[238] |
| ATTCTTTTCATAATCAAAATCAC | 8[447] | 8[425] |
| AATCGTTGAGTAACATTGGAATTACCTAATTACATTTAAC | 7[590] | 8[574] |
| ATTTTGCCAGAGGGGGTAATAGT | 8[279] | 8[257] |
| AGCGCCACCACGGAATACGCCTCAGACCAGAGCCACCACC | 7[422] | 8[406] |
| AAAAAAGGCAGCCTTTACAATCTTACCAGTTTG | 0[473] | 11[482] |
| TAATCGTAGCATTACCTGAGAGTCTG | 8[172] | 9[188] |
| CAAGTGCTGAGTAAGAAAATAAATCCTC | 6[405] | 2[392] |
| GGCTAAAGTACGGTGTCTGGAAG | 7[189] | 7[211] |
| CCTACATACGTAGCGGCCAGCCATTGCAACAGGTTTTT | 8[678] | 9[698] |
| CTATTTCGGAACGAGTGAGAATA | 4[377] | 4[355] |
| TCAACATCAGTTAAATAGCGAGAGTGAGACGACGATAAAA | 4[270] | 7[272] |
| AATAACGCGCGGGGAGAGG | 10[247] | 11[260] |
| AAGAGATTCATTTTGTTTAAGAGGAAGC | 6[237] | 2[224] |
| CAAATGGTTCAGAAGAACGAGTAGAT | 8[214] | 9[230] |
| AAAAGGGCGACAATTATTTATCC | 3[434] | 3[456] |
| ATAGCTGTTTCCTGGAACGTCCATAACGCCGTAAA | 11[63] | 6[70] |
| TGTAGGGGATTTAGTAACACTGAGTTTC | 2[349] | 3[349] |
| AAAAATCTACGTGCGTTTTAATT | 0[244] | 0[222] |
| AGAGTTTATACCAGTAGCACCTGAAACCATCGATA | 5[413] | 4[420] |
| GTGTATTAAGAGGCTGAGACTCC | 7[357] | 7[379] |
| GAAGTCAACCCAAATGGCAAAAGAATACTCGGAACAGAATCC | 9[273] | 5[286] |
| CGGTTAACAAAGCTGCTGTAACAACAAGGACGTTGGGAAG | 11[261] | 0[245] |
| ACTACCTTTAAACGGGTAACAGGGAGACGGGCA | 0[305] | 11[314] |
| AATCCAAAAAAAAAGGCTCCAAAA | 7[315] | 7[337] |
| GAGAGCCTCAGAACCGCATTTTCTGTAACGATCTAAAGTT | 11[345] | 0[329] |
| AAATCCCCGAAACAATTCATGAGGAAGT | 6[321] | 2[308] |
| TACCTAATATCAAAATCATTCAATATTACGTGA | 0[557] | 11[566] |
| GTATACAGGTAATGTGTAGGTAGTCAAATCACCAT | 5[161] | 4[168] |
| AACGTTGTAGAAACAGCGGATAGTTGGGCGGTTGT | 5[77] | 4[84] |
| GTTTATGTCACATGGGAATCCAC | 3[415] | 10[416] |
| ATATTCACAAACAAATTCATATG | 3[392] | 3[414] |
| GACCGGAAGCAATTGCGGGAGAA | 0[202] | 0[180] |
| TCAAGCAGAACCACCACTCACTCAGGTAGCCCGGAATAGG | 7[380] | 8[364] |
| AGCCTCCCCAGGGTCCGGCAAACGCG | 8[88] | 9[104] |
| TTCATTTTCTGCTAAACAACTGAACAACTAAAGGA | 5[329] | 4[336] |

| | | |
|---|---|---|
| TCGTTCACCGCCTGGCCCT | 10[331] | 11[344] |
| CGGAAGCACGCAAACTTATTAGCGTT | 8[424] | 9[440] |
| GAGCAAGGTGGCATTTACTCCAACAGGTTCTTTACGTCAACA | 9[189] | 5[202] |
| ATTGCGAATAATGTACAACGGAG | 4[335] | 4[313] |
| CTTTTTTTCGTCTCGTCGCTGGC | 8[111] | 8[89] |
| GACCGTCGAACGGGGAAGCTAATGCAGA | 6[531] | 2[518] |
| GCGTCATACATGCCCTCATAGTT | 0[370] | 0[348] |
| GAAAGTTCAACAATCAGCTTGCTTAGCTTTAATTGTATCG | 4[354] | 7[356] |
| TGTAAATCATGCTCCTTTTGATAATTGCTGAATAT | 5[203] | 4[210] |
| TTCACCTAGCGTGGCGGGTGAAGGGATACCAGTGCATAAAAA | 9[63] | 5[76] |
| ATTTGCCAAGCGGAACTGACCAACGAGTCAATCATAAGGG | 4[312] | 7[314] |
| TAGAACCTACCAGTCTGAGAGAC | 0[580] | 0[558] |
| GGGTTACCTGCAGCCAGCGGTGTTTTT | 4[51] | 4[29] |
| GAATTATCCAATAACGATAGCTTAGATT | 2[559] | 3[559] |
| TTGTCGTCTTTCTACGTAATGCC | 0[328] | 0[306] |
| ACTACTTAGCCGGAACGAGGCGC | 7[273] | 7[295] |
| TTTTTGTCCATCACGCAAATTCCGAGTAAAAGAGTCTTTTTT | 4[702] | 5[702] |
| TTTTTCGGGAGCTAAACAGGTTGTTAGAATCAGAGTTTTT | 0[694] | 1[694] |
| AATCATAATAACCCGGCGTCAAAAATGA | 6[489] | 2[476] |
| AGCAAGCCGTTTAAGAATTGAGT | 4[503] | 4[481] |
| AACAGAGTGCCTGGGGTTTTGCTCACAGAAGGATTAGGAT | 4[396] | 7[398] |
| CCAGCCAAACTTCTGATTGCCGTTTTGGGTAAAGTTAAAC | 4[102] | 7[104] |
| TGAAATTGTTTCAGGGAACTACAACGCC | 6[363] | 2[350] |
| GCCCGCACAGGCGGCCTTTAGTG | 7[63] | 7[85] |
| CAGTAAGAACCTTGAGCCTGTTTAGT | 8[550] | 9[566] |
| ACCAAATTACCAGGTCATAGCCCCGAGTTTTCATCGGCAT | 4[438] | 7[440] |
| TCTTATACTCAGAAAGGCTTTTGATGATATTGACACGCTATT | 9[357] | 5[370] |
| GCCTTATACCCTGTAATACCAATTCTTGCGCTC | 0[179] | 11[188] |
| TTTTTGCGTCCGTGCCTGCATCAGACGTTTTT | 9[25] | 6[21] |
| TTATGGCCTGAGCACCTCAGAGCATAAA | 2[181] | 3[181] |
| CGAGCACAGACTTCAAATACCTCAAAAGCTGCA | 0[221] | 11[230] |
| GCATCAAAAAGAAGTAAATTGGG | 3[224] | 3[246] |
| TAAGTAGAAGAACTCAAACTATCG | 7[651] | 7[673] |
| ATTTGGCAAATCAACAGTTGAAA | 7[609] | 7[631] |
| GTTGAAACAAACATCAAGAAAAC | 8[615] | 8[593] |
| GAATTGTAGCCAGAATGGATCAGAGCAAATCCT | 0[389] | 11[398] |
| GCTTGACCATTAGATACATTTCG | 8[237] | 8[215] |
| CTGAAAACCTGTTTATCAAACATGTAACGTCAA | 0[515] | 11[524] |

| | | |
|---|---|---|
| GACTTTCTCCGTGGCGCGGTTG | 0[76] | 0[54] |
| ACACAACATACGAGGGATGTGGCTATTAATCGGCC | 11[105] | 6[112] |
| TTTTTAACAATATTACCGTCGCTGGTAATATCCAGTTTTT | 6[694] | 7[694] |
| TGCCTGAACAGCAAATGAATGCGCGAACT | 6[657] | 2[644] |
| CAAATATCAAACCAGATGAATAT | 4[629] | 4[607] |
| CAATATGATATTGATGGGCGCAT | 4[167] | 4[145] |
| TTCTGGAATAATCCTGATTTTGCCCGGCCGTAA | 0[599] | 11[608] |
| TTAACAAGAGAATCGATGAACGG | 8[195] | 8[173] |
| GGGCCGGAAGCATAAAGTG | 10[121] | 11[134] |
| GTTTGAGGGGACCTCATTTGCCG | 4[125] | 4[103] |
| GTATTAGAGCCGTCAATAGATAA | 8[657] | 8[635] |
| GCTAATGCCGGAGAGGGTAGCTA | 7[147] | 7[169] |
| TACTTCTTTGATAAAAATCTAAA | 4[671] | 4[649] |
| GAAAGATCGCACTCCAGCCAGCT | 7[105] | 7[127] |
| TCAGGCTGCGCAACTGTTGGGAA | 8[153] | 8[131] |
| ATACCCTTCGTGCCACGCTGAACCTTGCTGAACCT | 5[623] | 4[630] |
| CATAATATTCCGTAATGGGATCCGTGCATCTGCCA | 5[119] | 4[126] |
| TTTTTATCCAATAAATCTCTACCCCGGTAAAACTAGCATG | 7[170] | 8[154] |
| CCGATAATAAAAGGGACTTAACACCGCGAACCACCAGCAG | 11[639] | 0[623] |
| CATCAGCGTCTGGCCTTCCACAGGAACCTGGGG | 0[137] | 11[146] |
| GGAATAACAGAGATAGACATACAAACTTGAGGATTTAGAA | 7[632] | 8[616] |
| CCGGAAGACGTACAGCGCCGCGATTACAATTCC | 0[95] | 11[104] |
| TTCGCGGATTGATTGCTCATTTTTTAAC | 2[139] | 3[139] |
| TAAAGGATTGTATAAGCGCACAAACGACATTAAATGTGAG | 11[135] | 0[119] |
| GATAAAAATTTTTAGCCAGCTTT | 0[160] | 0[138] |
| GATAGTGCAACATGATATTTTTGAATGG | 2[643] | 3[643] |
| GGATAACCTCACAATTTTTGTTA | 3[98] | 3[120] |
| TCAATAATAAAGTGTATCATCATATTCC | 2[601] | 3[601] |
| CAATAGGAACGCAAATTAAGCAA | 3[140] | 3[162] |
| GCGAAAGACGCAAAGCCGCCACGGGAAC | 2[97] | 3[97] |
| TTCCGAATTGTAAACGTGTCGCCAGCATCGGTGCGGGCCT | 7[128] | 8[112] |
| ACATCATTTAAATTGCGTAGAAACAGTACCTTTTA | 5[581] | 4[588] |
| AAGATAAAACAGTTGGATTATAC | 0[622] | 0[600] |
| AACACCCTAAAGGGAGCCC | 10[625] | 11[638] |
| GCATCGAGCCAGATATCTTTAGGACCTGAGGAAGGTTATC | 4[648] | 7[650] |
| CGTAAAGGTCACGAAACCAGGCAATAGCACCGCTTCTGGT | 4[144] | 7[146] |
| CGAGTAACAACCGTTTACCAGTC | 0[118] | 0[96] |
| GCCTTACGCTGCGCGTAAAATTATTTTTTGACGCTCAATC | 7[674] | 8[658] |

| | | |
|---|---|---|
| CCGAACCCCCTAAAACATCGACCAGTTTAGAGC | 0[641] | 11[650] |
| TGCGTACTAATAGTAGTTGAAATGCATATTTCAACGCAAG | 11[177] | 0[161] |
| GATTTTAGACAGGCATTAAAAATA | 0[664] | 0[642] |
| TGATTATCAGATATACGTGGCAC | 3[602] | 3[624] |
| TGGCAAGTTTTTTGGGGTC | 10[583] | 11[596] |
| TCAGCTAACTCACATTAAT | 10[163] | 11[176] |
| CTATTAGTCTTTCGCCGCTACAG | 3[644] | 3[666] |
| AACGCCAAAAGGCGGATGGCTTA | 4[251] | 4[229] |
| AAGAAACAATGACCGGAAACGTC | 4[461] | 4[439] |
| GTACATCGACATCGTTAACGGCA | 4[83] | 4[61] |
| ATACCACCATCAGTGAGGCCAAACCGTTGTAGCAA | 5[665] | 4[672] |

| **Biotinylated staple strands** | **5'-end** | **3'-end** |
|---|---|---|
| AACGCCAAAAGGCGGATGGCTTA | 4[251] | 4[229] |
| AAGAAACAATGACCGGAAACGTC | 4[461] | 4[439] |
| GTACATCGACATCGTTAACGGCA | 4[83] | 4[61] |
| ATACCACCATCAGTGAGGCCAAACCGTTGTAGCAA | 5[665] | 4[672] |
